# Phosphoinositide- and Collybistin-Dependent Synaptic Clustering of Gephyrin

**DOI:** 10.1101/2025.01.20.633899

**Authors:** Nele Burdina, Filip Liebsch, Arthur Macha, Joaquín Lucas Ortuño Gil, Pia Frommelt, Irina Rais, Fabian Basler, Simon Pöpsel, Guenter Schwarz

## Abstract

Gephyrin is the main scaffolding protein at inhibitory synapses clustering glycine and GABA type A receptors. At specific GABAergic synapses, the nucleotide exchange factor collybistin recruits gephyrin to the postsynaptic membrane via interaction with phosphoinositides. However, the molecular mechanisms underlying the formation, maintenance and regulation of collybistin-dependent gephyrin clusters remain poorly understood.

This study sheds light on the molecular mechanism of gephyrin cluster formation based on gephyrin self-oligomerization induced by collybistin, leading to the formation of a high-molecular weight (>5 MDa) gephyrin-collybistin complex, which is regulated in two ways: First, plasma-membrane phosphoinositides promote complex formation demonstrating their critical role in membrane targeting and stabilization of gephyrin-collybistin clusters at postsynaptic sites. Second, gephyrin phosphorylation at Ser325 abolishes complex formation with collybistin thus impairing collybistin-dependent gephyrin clustering at GABAergic synapses. Collectively, our data demonstrates a molecular mechanism for synaptic clustering of gephyrin which involves collybistin- and phosphoinositide-dependent formation of high-molecular gephyrin oligomers.

## INTRODUCTION

Efficient synaptic transmission requires precise accumulation of neurotransmitter receptors at the postsynaptic membrane. Scaffolding proteins provide structural support for various postsynaptic components, including neurotransmitter receptors, adhesion molecules and other functional elements (Sheng & Kim, 2011). At inhibitory postsynapses gephyrin (Geph) serves as the major scaffolding protein, clustering glycine receptors (GlyR) and specific subtypes of γ-aminobutyric acid type A receptors (GABA_A_R) (Fritschy et al., 2008). In addition to gephyrińs role in neuronal scaffolding, it catalyzes the last two steps of the molybdenum cofactor (Moco) biosynthesis (Feng et al., 1998; Stallmeyer et al., 1999). Geph is essential for postnatal survival as Geph-deficient mice die within the first postnatal day due to a lack of synaptically localized inhibitory receptors and impaired molybdoenzyme activity (Feng et al., 1998). In humans, Geph dysfunction has a severe impact on neurotransmission and has been associated with various brain disorders including epilepsy, Dravet-like syndrome, epileptic encephalopathy, schizophrenia and autism spectrum disorder (Dejanovic et al., 2014, 2015; Lionel et al., 2013; Macha, Liebsch, et al., 2022).

Geph is composed of an N-terminal G-, a central C-, and a C-terminal E-domain. The isolated G-domain and E-domain form trimers and dimers, respectively (E. Y. Kim et al., 2006; Schwarz et al., 2001; Sola et al., 2001). Geph E-domain dimers provide binding sites for the receptors and together with the G-domain trimers contribute to the formation of higher order oligomers into the postsynaptic gephyrin scaffold (Herweg & Schwarz, 2012; Pennacchietti et al., 2017; Specht et al., 2013). Interestingly, previous *in vitro* studies using recombinant full-length Geph found trimeric Geph with monomeric E-domains due to an inhibitory role of the central C-domain (Bedet et al., 2006; Sander et al., 2013). Therefore, the molecular mechanisms triggering E-domain dimerization at postsynaptic sites remain elusive and are thought to involve dynamic regulation via post translational modifications (PTMs) and interacting proteins relieving the inhibitory effect of the C-domain (Bedet et al., 2006; Sander et al., 2013).

At GABAergic synapses, the brain specific GTP/GDP exchange factor collybistin (CB) is a critical interactor binding to the domain boundary of the Geph C- and E-domain (Figure 1A) (Harvey et al., 2004). Experiments with CB-deficient mice revealed that CB is essential for the formation and maintenance of α2-/ γ2-subunit-containing GABA_A_R clusters that recruit gephyrin in various brain regions including hippocampus, cerebellum and basolateral amygdala (Papadopoulos et al., 2007, 2008). There are several CB splice variants (CB1-CB3), that all contain a tandem Dbl homology-(DH) and a pleckstrin homology-(PH) domain but differ in their N- and C-termini as well as the presence of a N-terminal regulatory src homology 3-(SH3) domain (Harvey et al., 2004). The PH-domain of CB interacts with phosphoinositides (PIPs) and was shown to mediate Geph transport towards synaptic membranes via interaction with phosphatidylinositol-3-monophosphate (PI(3)P) located at early/ sorting endosomes (Papadopoulos et al., 2017).

**Figure 1.**
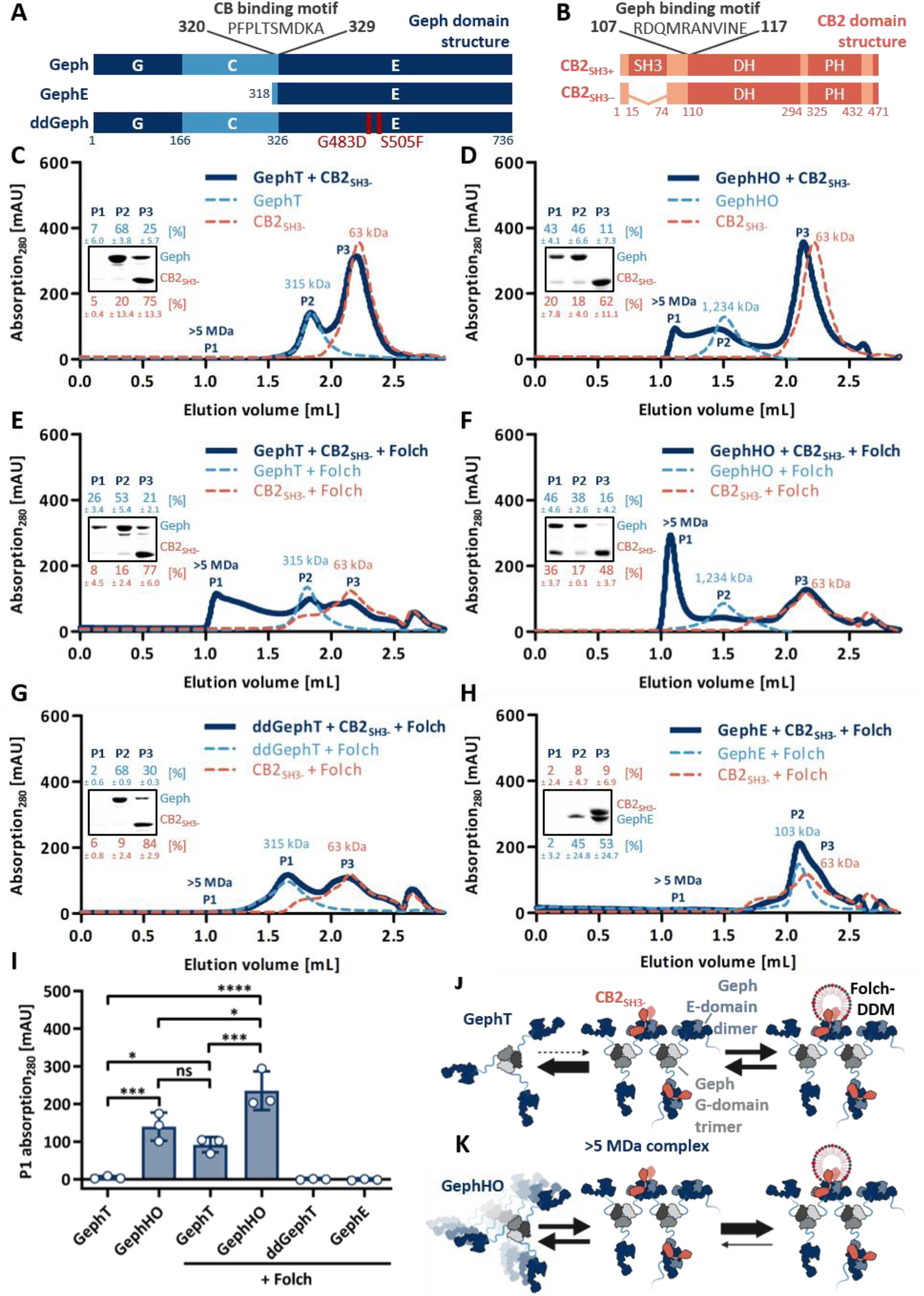
High-molecular weight Geph-CB2_SH3-_ complex formation is dependent on Geph self-oligomerization and promoted by Folch lipids. ***A***, Domain structure of the used Geph variants with the CB binding motif highlighted (Harvey et al., 2004). **B,** Domain structure of CB2 with and without SH3 domain. The Geph binding motif is highlighted (Grosskreutz et al., 2001; Tyagarajan, Ghosh, Harvey, et al., 2011). ***C – H*,** Representative SEC elution profiles of Geph variants mixed with CB2_SH3-_ at equimolar ratios (dark blue line), alone or in the presence of Folch. Single Geph (dashed line, light blue) and CB2_SH3-_ (dashed line, orange), with or without Folch, serve as a reference within each graph. The MWs of the single proteins as well as the formed complexes, determined according to standard protein calibration curve, are indicated. Insets depict SDS-PAGE analysis of peak 1 (P1), peak 2 (P2) and peak 3 (P3) of the respective Geph-CB2_SH3-_ interaction runs. Numbers above and below the representative SDS-PAGE image represent the mean relative band intensity ± SD [%] of Geph (light blue) and CB2_SH3-_ (orange), respectively, between P1, P2 and P3 (n = 3). ***I,*** Quantification of P1 absorption at 280 nm expressed as mean ± SD (n = 3). ddGephT and GephE are shown as a control depicting an absent Geph-CB2_SH3-_ compelx formation. The different oligomeric states of full-length WT-Geph were analyzed by 1way ANOVA (F(5,12)=37.89, *p*<0.0001, Bonferroni post-hoc test: GephT vs. GephHO *p*=0.0008 (***); GephT vs. GephT+Folch *p*=0.0299 (*)); GephT vs. GephHO+Folch *p*<0.0001 (****); GephHO vs. GephT+Folch *p*=0.7542 (ns); GephHO vs. GephHO+Folch *p*=0.0151 (*); GephT+Folch vs. GephHO+Folch *p*=0.0004 (***)). ***J, K,*** Proposed model for the formation of the >5 MDa Geph-CB2_SH3-_ compelx via Geph E-domain dimerization induced by CB2_SH3-_ and stabilized by Folch lipids comparing ***J,*** GephT and ***K,*** GephHO (created with BioRender.com).

CB isoform 2 containing the SH3 domain (CB2_SH3+_, Figure 1*B*) is the most abundantly expressed isoform within the brain (Harvey et al., 2004). CB isoforms containing the SH3-domain adopt a closed conformation impairing PIP binding and thereby preventing Geph recruitment to the membrane (Soykan et al., 2014; Xiang et al., 2006). Interaction partners binding to the PH-domain, such as the small GTPase TC10 or Neuroligin-2 (NL2), induce a conformational change, which allows PI(3)P binding (Kilisch et al., 2020; Mayer et al., 2013). Prolonged interaction with those interaction partners induces a phospholipid affinity switch towards plasma-membrane resident PIPs, such as PI(3,4)P_2_, PI(4,5)P_2_ and PI(3,4,5)P_3_, thereby enabling synaptic CB/Geph clustering (Kilisch et al., 2020; Poulopoulos et al., 2009; Schäfer et al., 2020). Impairments within CB-dependent synaptic Geph clustering lead to severe neurological dysfunctions such as epilepsy, anxiety, aggression and intellectual disabilities (Chiou et al., 2019; Kalscheuer et al., 2009). Despite the fundamental importance of CB-induced Geph cluster formation at GABAergic synapses, the molecular basis of formation, maintenance and regulation of Geph-CB clusters remains poorly understood.

Here, we characterized the molecular mechanism of Geph-CB complex formation by studying the oligomerization of Geph upon CB binding as well as the modulation of this interaction by PIPs and PTMs. We revealed that CB binding induces Geph self-oligomerization leading to the formation of a high-molecular weight (>5 MDa) Geph-CB complex, which was dependent on Geph E-domain dimerization. Our data showed that Geph-CB complex formation can be modulated in two directions: While plasma-membrane resident PIPs present at the postsynaptic membrane stabilize the complex, Geph phosphorylation at Ser325 inhibits the complex formation.

## MATERIAL AND METHODS

### Expression constructs

His_6_-tagged rat Geph P1 (Gehling et al., 2024) and rat GephE (Belaidi & Schwarz, 2013) constructs in pQE80L were described previously and were used for recombinant *E. coli* expression and purification. The amino acid exchanges for the generation of ddGeph, ddGephE (G483D and S505F) and the phosphomimicking Geph variants (T324D, S325D) were generated by site-directed mutagenesis.

For Geph expression in HEK GPHN^-/-^ cells, the construct mScarlet-IRES-mEGFP-Geph in pcDNA™ 5/FRT/TO (Thermo Fischer) was cloned via overlap extension PCR followed by Gibson cloning using the following plasmids as templates: pAAV-hSyn-mScarlet (gift from Karl Deisseroth; Addgene plasmid #131001; http://n2t.net/addgene:131001; RRID:Addgene_131001); pAAV-EF1a-mCherry-IRES-Flpo (gift from Karl Deisseroth; Addgene plasmid #55634; http://n2t.net/addgene:55634; RRID:Addgene_55634); pAAV-hSyn-fDIO-mEGFP-Gphn_P1 (Addgene plasmid #194978; http://n2t.net/addgene:194978; RRID:Addgene_194978). The construct pAAV-hSyn-fDIO-mScarlet-Gphn_P1 used for virus production for neuronal expression of Geph, was described previously (Addgene plasmid # 194972; http://n2t.net/addgene:194972; RRID:Addgene_194972). S325D-Geph in pQE80L was used as a template to generate mScarlet-IRES-mEGFP-S325D-Geph in pcDNA™ 5/FRT/TO and pAAV-hSyn-fDIO-mScarlet-Geph-S325D via Gibson assembly.

Rat CB2_SH3-_ in mCherry-C3 described previously (Dejanovic et al., 2015) was subcloned into pQE80L (Qiagen) by PCR using the restriction sites BamHI and SalI and into pEGFP-N1 including a moxBFP tag (gift from Erik Snapp; Addgene plasmid # 68064; http://n2t.net/addgene:68064; RRID:Addgene_68064) via Gibson cloning.

The *E. coli* expression constructs of LS-αβICD in pETDuet1 (Macha, Grünewald, et al., 2022) and Intein tagged GlyR β-loop in pTYB2 (Schrader et al., 2004) were described previously.

### Protein Purification

Recombinant His_6_-tagged full-length Geph variants (Geph, ddGeph, T324D-Geph, S325D-Geph) were expressed in *E. coil* BL21 Rosetta cells for 16 h at 18 °C while His_6_-tagged GephE and ddGephE were expressed for 16 h at 30 °C. Cells were harvested by centrifugation (5,000g, 10 min). The cell pellet was resuspended in lysis buffer (100 mM Tris/HCl pH 7.5, 250 mM NaCl) supplemented with 0.05 % (v/v) Tween-20, lysozyme (Sigma), protease inhibitors (cOmplete EDTA-free, Roche) and DNAse I (Roche) and stored at −20 °C until further use. All full-length Geph and GephE variants were affinity-purified using nickel-nitrilotriacetic acid resin (Ni-NTA, Cube Biotech) followed by preparative SEC as described in the following: Cells were lysed by two cycles of sonication (3 min, 30 s pulse, 30 s pause, 40 % amplitude) and pressure lysis (1,000 – 1,500 bar, EmulsiFlex high-pressure homogenizer, Avastin). After pelleting cell debris (50,000g, 1 h), the supernatant was incubated with Ni-NTA beads. Unbound proteins were removed by several washing steps using lysis buffer supplemented with 10 mM imidazole followed by lysis buffer supplemented with 20 mM imidazole. The protein was eluted in elution buffer (100 mM Tris/HCl pH 7.5, 250 mM NaCl, 400 mM imidazole). To further purify the protein and to separate the specific oligomeric states of Geph, preparative SEC was performed using a Superdex 200 16/60 column (120 ml column volume, GE Healthcare) equilibrated with storage buffer (25 mM Tris/HCl pH 7.5, 250 mM NaCl, 5 mM β-mercaptoethanol, 5 % (v/v) glycerol).

Recombinant His_6_-tagged CB2_SH3-_ was expressed in *E. coil* BL21 Rosetta cells for 16 h at 18 °C. Cell harvesting, NiNTA affinity purification as well as preparative SEC were carried out as described for Geph, except modified buffer compositions (Lysis buffer: 100 mM Hepes pH 8.0, 500 mM NaCl, 10 % (v/v) glycerol, 1 % (w/v) CHAPS, 1 mM β-mercaptoethanol; Elution buffer: 100 mM Hepes pH 8.0, 500 mM NaCl, 10 % (v/v) glycerol, 50 mM arginine, 50 mM glutamate, 300 mM imidazole, 1 mM β-mercaptoethanol; Storage buffer: 25 mM Tris/HCl pH 7.5, 250 mM NaCl, 10 % glycerol, 10 mM EDTA, 5 mM β-mercaptoethanol).

Recombinant Intein tagged GlyR β-loop was expressed in *E. coli* ER2566 for 16 h at 18°C and the affinity-purification was carried out using the IMPACT protein purification system (New England Biolabs) according to a previously described protocol (Schrader et al., 2004). After affinity purification the GlyR β-loop was further purified by preparative SEC using a Superdex 200 16/60 column (120 ml column volume, GE Healthcare) equilibrated with storage buffer.

Recombinant LS-αβICD, carrying a His_6_-tag at the LS-αICD subunit, was expressed and purified using Ni-NTA affinity purification followed by preparative SEC using storage buffer as previously described (Macha, Grünewald, et al., 2022).

All proteins were flash frozen in liquid nitrogen and stored at – 80°C until further use.

### Detergents and Lipids

Dodecyl-β-D-maltosid (DDM) was purchased from GLYCON Biochemicals GmbH, 1-palmitoyl-2-oleoyl-sn-glycero-3-phosphocholine (POPC), Folch lipids (Type I, Folch Fraction I) from Sigma and PIPs (PI(3)P, PI(4)P, PI(3,4)P_2_, PI(3,5)P_2_, PI(4,5)P_2_, PI(3,4,5)P_3_, C_16_ derivatives) from MoBiTec GmbH.

Stock solutions of POPC and Folch were prepared in chloroform, whereas the PIPs were dissolved in a mixture of chloroform, methanol and water according to the manufacturer’s instructions. POPC and PIP stock solutions were mixed and added to chloroform to obtain the desired lipid molar ratios of 10 mol % of the different phosphoinositides in POPC. The organic solvents were evaporated with a gentle argon stream. Lipid films were further dried under vacuum overnight. The lipid film was dissolved in DDM buffer (25 mM Tris/HCl pH 7.5, 250 mM NaCl, 1 % (w/v) DDM) by three cycles of vortexing (1 min) followed by sonication (10 min, SONOREX RK100H, Bandelin) to generate the desired lipid-DDM micelles (PIP-POPC-DDM micelles; Folch-DDM micelles).

### SEC interaction studies

The complex formation between the different Geph variants and CB2_SH3-_ as well as the self-oligomerization of Geph was analysed by analytical size SEC using a Superose 6 increase 5/10 column (3 mL, GE Healthcare). Therefore, the respective proteins were mixed at an equimolar ratio using 1.5 nmol each. For the conditions using lipids, additionally 1.5 nmol Folch lipids in DDM micelles or 1.5 nmol of the respective PIP-lipids in POPC-DDM micelles were added to the mixture. The samples were adjusted to a volume of 90 μL using storage buffer and afterwards incubated for 30 min on ice. After centrifugation (17 000g, 10 min, 4 °C) the samples were applied to the column and proteins were separated at 4°C and a flow rate of 0.2 mL/min in SEC buffer (25 mM Tris/HCl pH 7.5, 150 mM NaCl, 5 mM β-mercaptoethanol). For SEC runs in presence of lipid-DDM micelles, the SEC buffer was supplemented with 0.01 % DDM. Elution was monitored by monitoring absorbance at 280 nm. MWs were determined by comparison to the elution of standard proteins (Figure S2A). 100 µL fractions were collected and the desired peak fractions were further subjected to Coomassie stained SDS-Page analysis.

### ITC interaction studies

Isothermal titration Calorimetry (ITC) experiments were performed by titrating the GlyR β-loop (300 µM) into the respective Geph variants (30 µM) using a MicroCal Auto-ITC200 (Malvern). The used proteins were within the same batch of storage buffer. The ITC measurements were carried out at 37 °C using an injection volume of 1.25 – 1.5 μL, a spacing of 120 s between injections and an initial delay of 60 s. The reference power was set to 5 μCal/sec and the stirring speed to 750 rpm. Parameters and curves were fitted and calculated by the MicroCal Analysis software and Origin7.

### *In vitro* Moco assay

The *in vitro* Moco synthesis assay was performed as previously described (Belaidi & Schwarz, 2013). For the analysis of Moco production of Geph in complex with CB2_SH3-_, both proteins were mixed at an equimolar ratio at a concentration of 25 µM either in absence or presence of 25 µM Folch-lipids in DDM micelles. 300 pmol of the complex or the respective Geph variant alone were used for the assay. Samples without Geph (-Geph) or without molybdenum (-Mo) were used as control and show Moco produced either chemically or by trace amounts of molybdenum in the buffers.

### SDS-PAGE, Coomassie and Western Blot

Samples were supplemented with 1x sample buffer (5x: 250 mM Tris/HCl pH 6.8, 30 % glycerol, 0.1 % Bromophenol blue, 10 % SDS, 5 % β-mercaptoethanol) and incubated for 5 min at 95 °C. Protein separation was performed with 12% SDS acrylamide gels followed by Coomassie-staining (30 % EtOH, 10 % acetic acid, 0.25 % Coomassie brilliant blue R250) or immunoblotting using standard protocols with a chemiluminescence and an ECL system. The following antibodies were used and diluted in Tris-buffered saline/ 0.5 % Tween containing 1 % dry milk: anti-Geph E-domain (3B11, 1:10, self-made, RRID: AB_887719); anti-GAPDH (1:1000, G9545, Sigma), anti-mouse HRP-coupled (1:10 000, AP181P, Sigma); anti-rabbit HRP-coupled (1:10 000, AP187P, Sigma). Image acquisition of coomassie stained Gels and immunoblot detection were performed with a ChemiDocTM Imaging System (BioRad). Band intensities were quantified using Image Lab 6.1 (BioRad).

### Generation of HEK GPHN^−/−^ cells

Human embryonic kidney (HEK293) cells were cultured in Dulbecco’s modified Eagle’s medium supplemented with 10 % FCS and 2 mM L-glutamine at 37°C and 5 % CO_2_. Geph-deficient HEK293 cells were generated using the double-nickase CRISPR/cas9 approach described previously (Ran et al., 2013). Two pX335 constructs (Addgene #42335), each encoding one *GPHN*-specific guide RNA (gRNA 1: TACTAACCACGACCATCAAA; gRNA 2: CTCAGGAATGTCCATTGGCC) were generated. HEK293 cells were transfected daily with both pX335 constructs for five days using GeneJuice transfection reagent (Novagen) according to manufactures protocol. After four days, cells were splitted 1:2. After five days of dilution. Therefore, cells were detached via trypsin/EDTA treatment, diluted to a concentration of 0.5 cells/100 μL and 100 μl were plated into 96-well plates. Single colonies were grown to ∼70 % confluency and transferred onto 12-well plates. HEK293 GPHN^−/−^ cells were identified via western blot.

### HEK GPHN^−/−^ transfection

HEK293 GPHN^−/−^ cells were plated onto collagen (PanBiotech) coated glass coverslips and transfected using 400 ng of the respective plasmid DNA with polyethylenimine (PEI, Sigma) according to the manufacturer’s protocol. 40 h after PEI transfection, cells were fixed.

### Virus preparation

Production of rAAV particles was carried out in HEK293 cells, followed by PEG/NaCl precipitation and chloroform extraction according to a previously described protocol (Kimura et al., 2019). Purity of the rAAVs was assessed using SDS-PAGE and titers were determined using Gel green® (Biotium) according to a previously described protocol (Xu et al., 2020). The fluorescence was detected at 507 ± 5 nm (excitation) and 528 ± 5 nm (emission) using a plate reader with monochromators (Tecan Spark).

### Primary neurons

Dissociated primary hippocampal cultures were prepared from C57BL/6NRj embryos (E17.5) of either sex. Therefore, pregnant dams were killed by cervical dislocation. E17.5 embryos were isolated and decapitated as quickly as possible. After dissociation, cells were seeded on poly-L-lysine coated cover slips in neurobasal medium supplemented with B-27, N-2, and L-glutamine (*Thermo Fisher Scientific*). After 9 d *in vitro* (DIV), neurons were transfected with 400 ng plasmid DNA using Lipofectamine 2000 (Life Technologies) according to the manufacturer’s protocol. After 10 DIV, neurons were infected with 1 x 10^8^ viral genome copies using rAAV particles diluted in neurobasal medium supplemented with B-27, N-2, and L-glutamine (*Thermo Fisher Scientific*). After 15 DIV, cells were fixed and used for Immunocytochemistry.

### Cell fixation and immunocytochemistry

All steps were performed at room temperature (RT). Cells were fixed using 4 % paraformaldehyde in PBS for 15 min, followed by three washing steps using 50 mM ammonium chloride in PBS. For immunocytochemistry, the cells were blocked/ permeabilized for 1 h using 10 % goat serum, 1 % BSA, 0.2 % Triton X-100 in PBS. Cells were incubated for 1 h with the following antibodies: anti-vesicular GABA transporter (vGAT) (1:1000, #131003) for inhibitory presynaptic terminals; anti-GABA_A_R γ2 (1:500, #224004) for postsynaptic GABA_A_Rs and subsequently washed using PBS. The following secondary antibodies were used: goat anti-rabbit AlexaFluor 488 (1:500, #A-11034, Thermo Fisher Scientific), and goat anti-guinea pig AlexaFluor 647 secondary antibodies (1:500, #ab150187, Abcam). The coverslips were mounted onto glass slides using a homemade Mowiol/Dabco solution, dried overnight at room temperature and stored at 4°C.

### Confocal microscopy and image analysis

Images were acquired with stacks of 3 μm z-step size, 2048 x 2048 (144.77 x 144.77 μm) using the Leica TCS SP8 LIGHNING upright confocal microscope with an HC PL APO CS2 63x/1.30 glycerol objective. The microscope was equipped with hybrid detectors (Leica HyD) and diode lasers with 405, 488, 522 and 638 nm. LIGHTNING adaptive deconvolution of the mounting medium “Mowiol” was used, which is capable of theoretical resolutions down to 120 nm and 200 nm lateral and axial, respectively. Images were segmented and analyzed in an automated fashion using ImageJ/FIJI and an adapted previously described macro (Liebsch et al., 2023).

### Subcellular fractionation of brain tissue

Fractionation of cytosolic and membrane associated gephyrin from pig brain tissue of either sex was performed as described in the following. In brief, frozen pig brain tissue was mortared into fine powder and dissolved in 10 tissue volumes tissue lysis buffer (20 mM Hepes pH 8.0, 150 mM NaCl, 5 % glycerol, 5 mM EDTA, 50 mM DTT, 20 mM NEM) supplemented with Protease inhibitor (cOmplete, Roche). Cells were lysed mechanically using a Potter with teflon pestle (1100 rpm, 30 strokes) followed by sonication (2x 10 s, 30 % amplitude). Cell debris and unbroken cells were removed by centrifugation at 4000xg (4 °C, 10 min) and the supernatant was used for ultracentrifugation (185.000xg, 1 h, 4 °C, Beckman Type 70.1 Ti rotor). The supernatant was used as the cytosol sample and was used directly for blue native page sample preparation. The pellet represents the membrane sample and was dissolved in tissue lysis buffer supplemented with 1 % (w/v) DDM and mechanically disrupted using a glass potter (50x). After an incubation time of 1 h (4°C, head over tail), solubilized membranes were centrifuged (185.000xg, 30min, 4°C, Beckman Type 70.1 Ti rotori) and the supernatant was used for blue native page sample preparation.

### Blue native PAGE analysis

Samples for blue native PAGE analysis were prepared by 1:3 dilution using H_2_O and the addition of 10x blue native loading buffer (312 mM imidazole, 500 mM 6-aminohexanoic acid, 5 % CBB G250, 6.25 mM EDTA, 40 % glycerol). Samples were applied to 4 – 16 % native PAGE (TM Novex ® Bis-Tris Gel) and protein separation was performed according to the manufacturers protocol. Afterwards gels were subjected to western blotting.

### Mass photometry measurements

Mass Photometry measurements were conducted using a Refeyn TwoMP mass photometer (Refeyn Ltd.). Standard microscopy glass slides were cleaned with MilliQ water and isopropanol. Residual lint was removed by a stream of compressed air, and 3×2-well silicon sample well cassettes were attached to the clean glass slides. The detector was cleaned with isopropanol, and the coverslips were mounted onto the detector using one drop of immersion oil. For each mass photometry measurement, the protein samples were pre-diluted to approximately 100 nM using dilution buffer (25 mM Tris/HCl pH 7.5, 150 mM NaCl, 5 mM β-mercaptoethanol, 1 % (v/v) glycerol). 19 µl of the dilution buffer were placed into one silicon well and the focal plane was automatically estimated via the droplet dilution feature in the chamber, resulting in a final concentration of 5 nM. The measurement was conducted for 60 s, after which the contrast histograms were analyzed in the software DiscoverMP (Refeyn Ltd.) using a previously acquired mass calibration with bovine serum albumin and human transglutaminase 2. Average masses were estimated using a standard gaussian fitting over the respective peaks. For each measurement, an average of 3,000 counts were targeted as a quality control measure to produce comparable results, and the sample concentrations were adjusted accordingly.

### Statistical analysis

Individual data points, mean, standard deviation and confidence intervals (CIs) are displayed in the figures. The used statistical tests are indicated in the figure legends. Visualization and statistical analysis were performed with GraphPad Prism 7 and R (version 4.3.2.) using the following packages: Dabestr 0.3.0, Pacman 0.5.1, rio 1.0.1, tidyverse 2.0.0, ggrepel 0.9.5, RColorBrewer 1.1-3, svglite 2.1.3, ggpubr 0.6.0, rstatix 0.7.2, effsize 0.8.1. Data was tested for normality, using Shapiro–Wilk test with a violation limit of p < 0.01. Datasets that were not normally distributed were analyzed using nonparametric tests. Datasets that were normally distributed were analyzed with the indicated parametric tests. Statistical significance is designated as \**p* < 0.05, \*\**p* < 0.01, \*\*\**p* < 0.001 and \*\*\*\**p* < 0.0001. For each experiment, the number of cells (n) and biological/ technical replicates (n) are reported within the figure legends.

### Ethics statement

All relevant ethical regulations for animal testing and research were followed, and the experiments were authorized by the local research ethics committees (Germany, Landesamt für Natur, Umwelt und Verbraucherschutz Nordrhein-Westfalen, reference 2021.A450).

## RESULTS

### Geph-CB2_SH3-_ complex formation requires Geph oligomerization

Geph E-domain dimerization is essential for synaptic Geph clustering (Saiyed et al., 2007). However, within full-length gephyrin, E-domain dimerization is inhibited by the C-domain (Bedet et al., 2006; Sander et al., 2013). We probed whether binding events within the Geph C-domain, such as the interaction with CB (Harvey et al., 2004), could relieve this inhibitory effect and thereby induce Geph oligomerization. To study the oligomerization of Geph upon CB binding, size exclusion chromatography (SEC) based interaction studies using recombinantly expressed and purified proteins were performed (Figure 1). SEC, Dynamic light scattering, chemical cross-linking and atomic force microscopy studies revealed that recombinant full-length Geph expressed in *E. coli* forms stable trimers alongside with higher oligomeric states (Saiyed et al., 2007; Sander et al., 2013; Schrader et al., 2004; Sola et al., 2004). Therefore, the different oligomeric states of *E. coli* expressed and affinity purified Geph were separated via SEC (Figure S1A) resulting in Geph trimers (GephT, 1.83 mL = 315 kDa, Figure 1C) and Geph high oligomers (GephHO, 1.50 mL = 1,234 kDa, Figure 1D). To study the complex formation between the different oligomeric states of recombinant Geph and the constitutively active (open conformation) CB2 splice variant lacking the SH3-domain (CB2_SH3-_, Figure 1B) (Harvey et al., 2004), equimolar amounts of both proteins were coincubated (30 min) and subsequently analyzed by SEC (Figures 1C and 1D). Surprisingly, in case of GephT no complex formation with CB2_SH3-_ was observed (Figure 1C). In contrast, GephHO formed a high-molecular weight complex with CB2_SH3-_ eluting at the void volume of the SEC column, corresponding to a molecular weight (MW) larger than 5 MDa based on the fractionation limit of the resin (Figure 1D).

Blue-native PAGE analysis of Geph from pig brain revealed the presence of cytosolic Geph trimers and membrane-associated higher oligomers (Figure S1B). The observed band height of cytosolic Geph from brain extracts corresponded to the size of recombinant GephT while the band height of membrane-associated Geph corresponded to the size of recombinant GephHO (Figure S1B). Thus, our data suggests that trimeric Geph is mainly found within the cytosol, while the formation of distinct larger Geph oligomers is induced at sub-membranous sites. In conclusion, Geph-CB2_SH3-_ complex formation depends on the oligomeric state of Geph and is promoted for GephHO, found at sub-membranous sites, compared to GephT, mainly found within the cytosol.

### Lipids stabilize the Geph-CB2_SH3-_ complex

The ability of CB to bind PIPs is crucial for Geph trafficking towards the synaptic membrane (Harvey et al., 2004; Papadopoulos et al., 2017). To test the effect of lipids on the high-molecular weight Geph-CB2_SH3-_ complex formation, we performed similar SEC experiments in the presence of bovine brain lipid extract Folch fraction I (from here on referred to as Folch) (Eggers & Schwudke, 2016).

For the SEC studies, Folch lipids were solubilized in DDM detergent micelles and added to equimolar amounts of CB2_SH3-_ and the respective Geph multimers (Figures 1E and 1F). The addition of DDM alone did not alter the Geph-CB2_SH3-_ complex formation in case of both oligomeric states compared to the condition without any detergent (Figures S2E and S2F). As expected, CB2_SH3-_ alone did interact with the Folch-DDM micelles, as a peak shift was detected when compared to the conditions without Folch (Figure S2B). The elution profile of GephT and GephHO was not altered by the addition of Folch lipids, indicating that the lipids did not affect the oligomerization of Geph (Figures S2C and S2D).

The addition of Folch lipids promoted complex formation with CB2_SH3-_ for both oligomeric states of Geph (Figures 1E and 1F). In case of GephHO the addition of Folch lipids induced a significant increase of the void volume peak (>5 MDa, P1, Figure 1F) compared to the condition without lipids (Figure 1D and 1I, *p* = 0.0151). Surprisingly, in the presence of Folch lipids also GephT formed a complex with CB2_SH3-_ eluting at the void volume of the column (>5 MDa, P1, Figure 1E), while in the absence of lipids no peak shift was detected (Figures 1C and 1I, *p* = 0.0299). The height of the void volume peak P1 was significantly smaller in case of GephT (Figure 1E) compared to the complex formed with GephHO (Figures 1F and 1I, *p* = 0.0004), indicating that also in the presence of Folch lipids the Geph-CB2_SH3-_ complex formation is less efficient for GephT compared to GephHO. In summary, lipid binding to CB strengthened the interaction with Geph, enabling complex formation with both oligomeric states of Geph.

Previous studies mapping the binding sites of Geph and CB suggest that one Geph monomer binds to one CB2_SH3-_ monomer (Grosskreutz et al., 2001; Harvey et al., 2004). Thus, GephT (315 kDa, Figure 1C) harbors three potential binding sites for CB2_SH3-_ (63 kDa, Figures 1C – 1F) while GephHO (1,234 kDa, Figure 1D), containing approximately four Geph trimers, harbors twelve potential binding sites. Based on the MWs determined by comparison to the elution of standard proteins (Figure S2A), the resulting complexes would correspond to a MW of approximately 504 kDa for GephT and 1,990 kDa for GephHO. Thus, the formed >5 MDa Geph-CB2_SH3-_ complex exceeds the theoretical MW of a 1:1 complex for GephT (∼10-fold) and for GephHO (∼2.5-fold). We therefore conclude that either a conformational change was induced upon CB2_SH3-_ binding, leading to an extended hydrodynamic radius or additional Geph and/or CB2_SH3-_ subunits were associated to enlarge the Geph-CB2_SH3-_ complex.

### High-molecular weight Geph-CB2_SH3-_ complex is based on Geph self-oligomerization via E-domain dimerization

To address the question whether the formation of the high-molecular weight GephHO-CB2_SH3-_ complex is based on Geph oligomerization via Geph E-domain dimerization, we performed SEC interaction studies with oligomerization-deficient Geph variants. Therefore, an E-domain dimerization-deficient Geph variant (ddGeph) was generated by introducing two amino acid exchanges in the E-domain (G483D and S505F) both located outside the CB binding motif (Figure 1A). Krausze and colleagues (2016) identified specific mutations in the plant-ortholog of the protein (Krausze et al., 2016). Since single amino acid exchanges only partially shifted the equilibrium from dimerization to monomerization of Cnx1E (Krausze et al., 2016), we targeted both amino acid residues in Geph to achieve full E-domain monomerization.

G483D is located within an α-helix, that is directly involved in the dimerization interface while S505F is located within a structurally conserved β-sheet in close proximity to an α-helix within the dimerization interface (Figure S3A). To study the effect of the dual substitution on E-domain dimerization, we characterized the oligomerization behavior of the isolated Geph E-domain carrying both substitutions (ddGephE). SEC experiments revealed that ddGephE elutes with a smaller hydrodynamic radius compared to the WT-Geph E-domain (GephE, Figure S3C), being consistent with a switch from the dimeric to the monomeric state. Additionally, mass photometry measurements confirmed that the measured MW of ddGephE (49 ± 4.3 kDa) correlates to a monomer (theoretical MW = 47 kDa) while the measured MW of WT-GephE (89 ± 5.5 kDa) matches to that of a dimer (theoretical MW = 94 kDa, Figures S3D and S3E).

SEC studies of full-length ddGeph directly after affinity purification revealed that besides trimer formation, no higher oligomeric states were observed (Figure S3B), indicating that E-domain dimerization is essential for the formation of GephHO. As WT-GephT formed a high-molecular weight complex with CB2_SH3-_ exclusively when Folch lipids were present (Figures 1C and 1E), we repeated those experiments in the presence of Folch lipids now with trimeric ddGeph (ddGephT). Interestingly, ddGephT did not form a complex together with CB2_SH3-_ and Folch lipids (Figure 1G). This finding further supports our hypothesis, that the high-molecular weight Geph-CB2_SH3-_ complex is based on a Geph oligomerization that requires E-domain dimerization (Figures 1J and 1K).

The formation of a Geph scaffold additionally requires the trimerization of the G-domain (Saiyed et al., 2007). Therefore, we again performed SEC experiments with WT-GephE, which still contains the entire CB binding motif but lacks the trimerizing G-domain (Figure 1A). Again, no complex formation with CB2_SH3-_ and Folch lipids was detected (Figure 1H). This finding led us conclude, that also G-domain trimerization is required for the high-molecular weight complex formation.

In summary, our results using oligomerization-deficient Geph variants show that both Geph E-domain dimerization as well as G-domain trimerization are required for the high-molecular weight Geph-CB2_SH3-_ complex formation. Thus, we conclude that the formation of the high molecular weight Geph-CB2_SH3-_ complex is based on Geph self-oligomerization induced by interaction with CB2_SH3-_ (Figures 1J and 1K).

### Geph-CB2_SH3-_ complexes are functionally active

To confirm that the >5 MDa Geph-CB2_SH3-_ complex contains properly folded, active proteins, we performed a functional analysis of the complex (Figure S4). Geph catalyzes the last two steps of the Moco biosynthesis, with the G-domain catalyzing the adenylation of molybdopterin (MPT) and the E-domain promoting the incorporation of molybdate into activated MPT-AMP (Figure S4A) (Feng et al., 1998; Stallmeyer et al., 1999). We performed an *in vitro* assay (Belaidi & Schwarz, 2013), to measure Moco production by GephHO in complex with CB2_SH3-_. The resulting enzymatic activity was not impaired as no significant difference in Moco production compared to GephHO alone was observed. Also, the addition of Folch lipids did not alter Moco synthesis of GephHO within the complex (Figure S4B).

Next, we investigated whether the neuronal receptor binding function of Geph is preserved in the >5 MDa Geph-CB2_SH3-_ complex. The interaction between the inhibitory neurotransmitter receptors and Geph is facilitated through the intracellular cytosolic domains (ICDs) of the receptors (Kowalczyk et al., 2013; Meyer et al., 1995; Tretter et al., 2011). Therefore, we studied whether the Geph-CB2_SH3-_ complex can recruit binding peptides of inhibitory neurotransmitter receptors using a recently established pentameric soluble protein platform.(Macha, Grünewald, et al., 2022) The platform is based on the insertion of the full-length ICDs of the GlyR α- and β-subunits, into the soluble pentameric yeast lumazine synthase, resulting in the heteropentameric protein LS-αβICD (Figure S4C). As the GlyR and GABA_A_R compete for the same binding pocket within the interface of the dimerized Geph E-domain (Maric et al., 2014), the GlyR ICDs represent a suitable model to study the receptor binding ability.

We performed SEC experiments to investigate whether LS-αβICD is recruited to the >5 MDa GephHO-Folch-CB2_SH3-_ complex (Figure S4D). Indeed, the peak corresponding to LS-αβICD (P3, 176 kDa, dashed grey line) was reduced while the dominant void volume peak (P1) increased, indicating that LS-αβICD interacts with the >5 MDa GephHO-Folch-CB2_SH3-_ complex (P1, bold dark blue line, Figure S4D). In contrast, LS-αβICD with GephHO alone did only form a smaller complex of 1,397 kDa in size (P2, bold light blue line, Figure S4D) while no interaction was detected between LS-αβICD and CB2_SH3-_(bold orange line, Figure S4D).

The LS-αβICD-GephHO-CB2_SH3-_ complex formation was further confirmed via SDS-Page analysis of corresponding peak fractions (Figure S4E). The band ratio of LS-αβICD within the void volume peak P1 was increased to 63 % for the LS-αβICD-GephHO-CB2_SH3-_ complex compared to 20 % and 11 % in case of LS-αβICD with GephHO alone, respectively. These results further support the conclusion that LS-αβICD was recruited by the >5MDa GephHO-CB2_SH3-_ complex. In summary, Geph within the >5 MDa Geph-CB2_SH3-_ complex remained enzymatically active and is able to recruit inhibitory neurotransmitter receptor binding sites, which is essential for the proper function of CB-Geph clusters at inhibitory postsynapses.

### PIPs, present at the plasma membrane, stabilize the Geph-CB2_SH3-_ complex

Next, we aimed to determine which types of lipids promote Geph-CB2_SH3-_ complex formation. CB is known to interact with PIPs via its PH domain with a high specificity for PI(3)P, while interactors like TC10 or NL-2 induce a phospholipid affinity switch towards plasma membrane resident PIPs, such as PI(3,4)P_2_, PI(4,5)P_2_ and PI(3,4,5)P_3_ (Kilisch et al., 2020; Papadopoulos et al., 2017; Poulopoulos et al., 2009; Schäfer et al., 2020).

To study the effect of PIPs on the Geph-CB2_SH3-_ complex formation, we doped POPC DDM micelles with 10 mol % of different PIPs, namely PI(3)P, PI(4)P, PI(3,4)P_2_, PI(3,5)P_2_, PI(4,5)P_2_ and PI(3,4,5)P_3_ (Figure 2C). Our subsequent PIP ‘screen’ was performed with GephHO as it showed a more stable complex formation with CB2_SH3-_ compared to GephT (Figures 1C – 1F). To study the effect of the PIPs, the respective lipid-DDM micelles were added to equimolar amounts of CB2_SH3-_ and GephHO and complex formation was studied by SEC (Figure 2A). The addition of POPC (Figure 2A) did not alter the Geph-CB2_SH3-_ complex formation as compared to DDM only or no additive added (Figure S2F).

**Figure 2.**
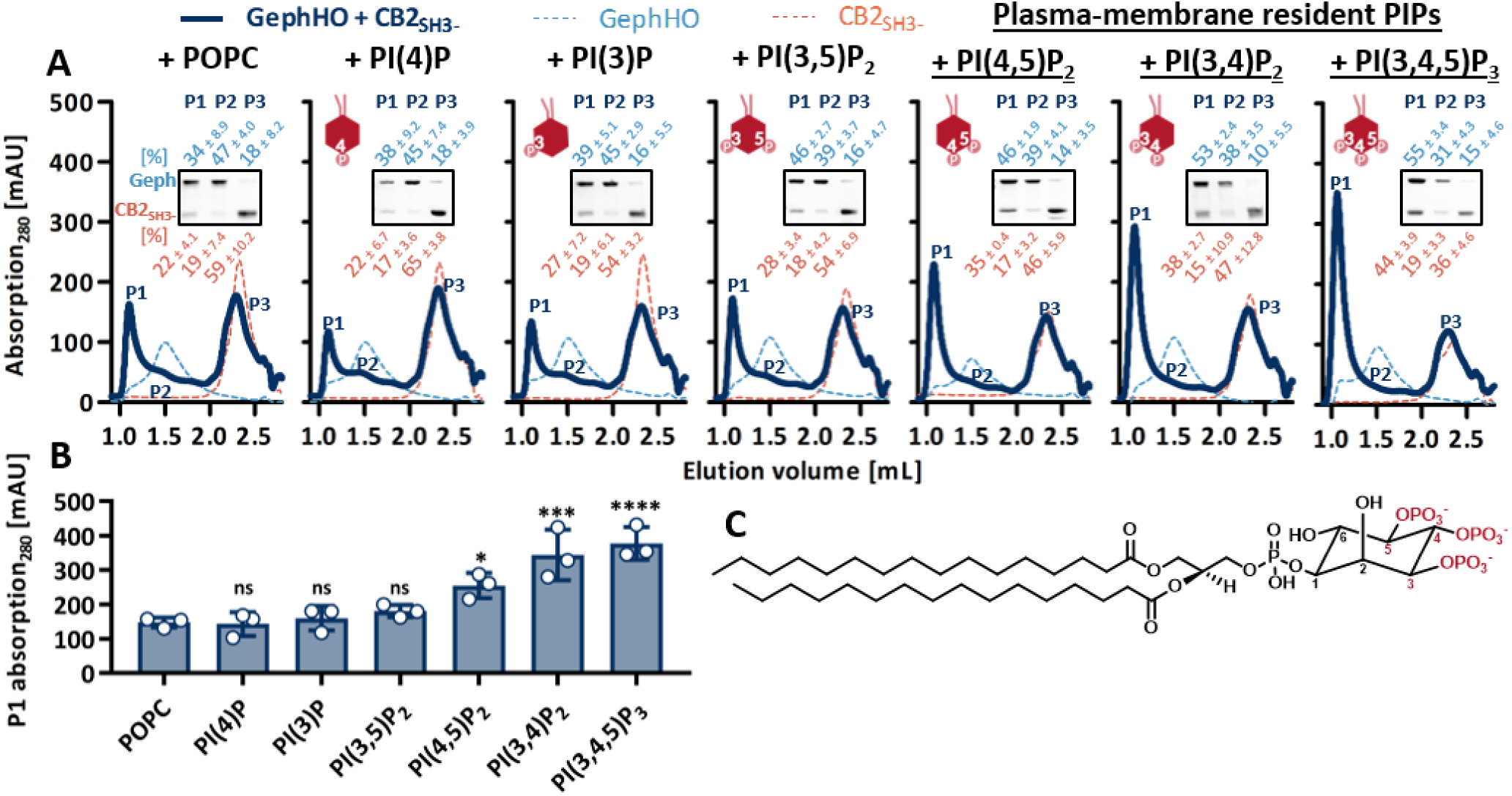
Plasma-membrane resident PIPs stabilize the high-molecular weight Geph-CB2_SH3-_ complex. ***A,*** Representative SEC elution profiles of GephHO mixed with CB2_SH3-_ at equimolar ratios, in the presence of the respective POPC-/ PIP-lipid DDM micelles (dark blue line). Single Geph (dashed line, light blue) and CB2_SH3-_ (dashed line, orange) together with the respective POPC-/ PIP-lipid DDM micelles serve as a reference within each graph. Insets depicting SDS-PAGE analysis of peak 1 (P1), peak 2 (P2) and peak 3 (P3) of the respective Geph-CB2_SH3-_ interaction run. Numbers above and below the representative SDS-PAGE image represent the mean relative band intensity ± SD [%] of Geph (light blue) and CB2_SH3-_ (orange), respectively, between P1, P2 and P3 (n = 3). Depiction of the respectively used PIP created with BioRender.com. ***B,*** Quantification of P1 absorption at 280 nm expressed as mean ± SD (n = 3). PIP treated conditions were compared to the POPC control condition, revealing that P1 absorption is significantly increased for plasma-membrane resident PIPs (1way ANOVA F(6,14)=16.64, *p*<0.0001, Dunnett’s post-hoc test: PI(4)P *p*=0.9998 (ns); PI(3)P *p*=0.9980 (ns); PI(3,5)P_2_ *p*=0.8317 (ns); PI(4,5)P_2_ *p*=0.0306 (*); PI(3,4)P_2_ *p*=0.0003 (***); PI(3,4,5)P_3_ *p*=0.0001 (****)). ***C,*** Molecular structure of a PIP with the possible phosphorylation sites at the inositol ring highlighted in red (created with ChemDraw).

The phosphatidylinositol monophosphates PI(3)P and PI(4)P as well as PI(3,5)P_2_ did not improve the Geph-CB2_SH3-_ complex formation as the height of the >5 MDa peak P1 did not increase compared to the POPC control (Figures 2A and 2B). Interestingly, only PIPs known to be present at the plasma membrane, PI(3,4)P_2_, PI(4,5)P_2_ and PI(3,4,5)P_3_ (Ueda, 2014), did significantly increase the height of P1 compared to the POPC control condition (Figures 2A and 2B). The presence of phosphatidylinositol triphosphate PI(3,4,5)P_3_ induced the strongest increase of P1 with 378 ± 39 mAU (*p* < 0.0001) followed by PI(3,4)P_2_ with 344 ± 60 mAU (*p* = 0.0003) and PI(4,5)P_2_ with 255 ± 30 mAU (*p* = 0.0306) compared to 148 ± 12 mAU for POPC (Figure 2B).

Changes in P1 height were further confirmed by SDS-Page analysis of the peak fractions, showing the respective increase of band intensity corresponding to Geph and CB2_SH3-_ within the >5 MDa complex compared to the control condition using POPC (insets Figure 2A). Again, the strongest increase was observed for PI(3,4,5)P_3_ with an increase of 21 % GephHO and 22 % CB2_SH3-_ within P1 compared to the POPC control. In summary, our SEC studies showed that PIPs present at the plasma membrane promote high-molecular weight Geph-CB2_SH3-_ complex formation, indicating that the Geph-CB scaffold is stabilized and maintained at synaptic membranes via the interaction with those PIPs.

### Phosphomimicking mutant S325D-Geph does not form a stable complex with CB2_SH3-_

Geph is subject to various PTMs such as phosphorylation, palmitoylation and nitrosylation that can affect the structure and scaffolding properties of Geph, its trafficking as well as its ability to interact with partner proteins (Tyagarajan, Ghosh, Yévenes, et al., 2011). There are two known phosphorylation sites within the CB binding motif of Geph, Thr324 and Ser325 (Herweg & Schwarz, 2012; Ogino et al., 2019). We studied the impact of both phosphorylations on the Geph-CB complex formation by exchanging the respective residues to aspartates resulting in the phosphomimicking variants T324D-Geph and S325D-Geph (Figure 3A).

**Figure 3.**
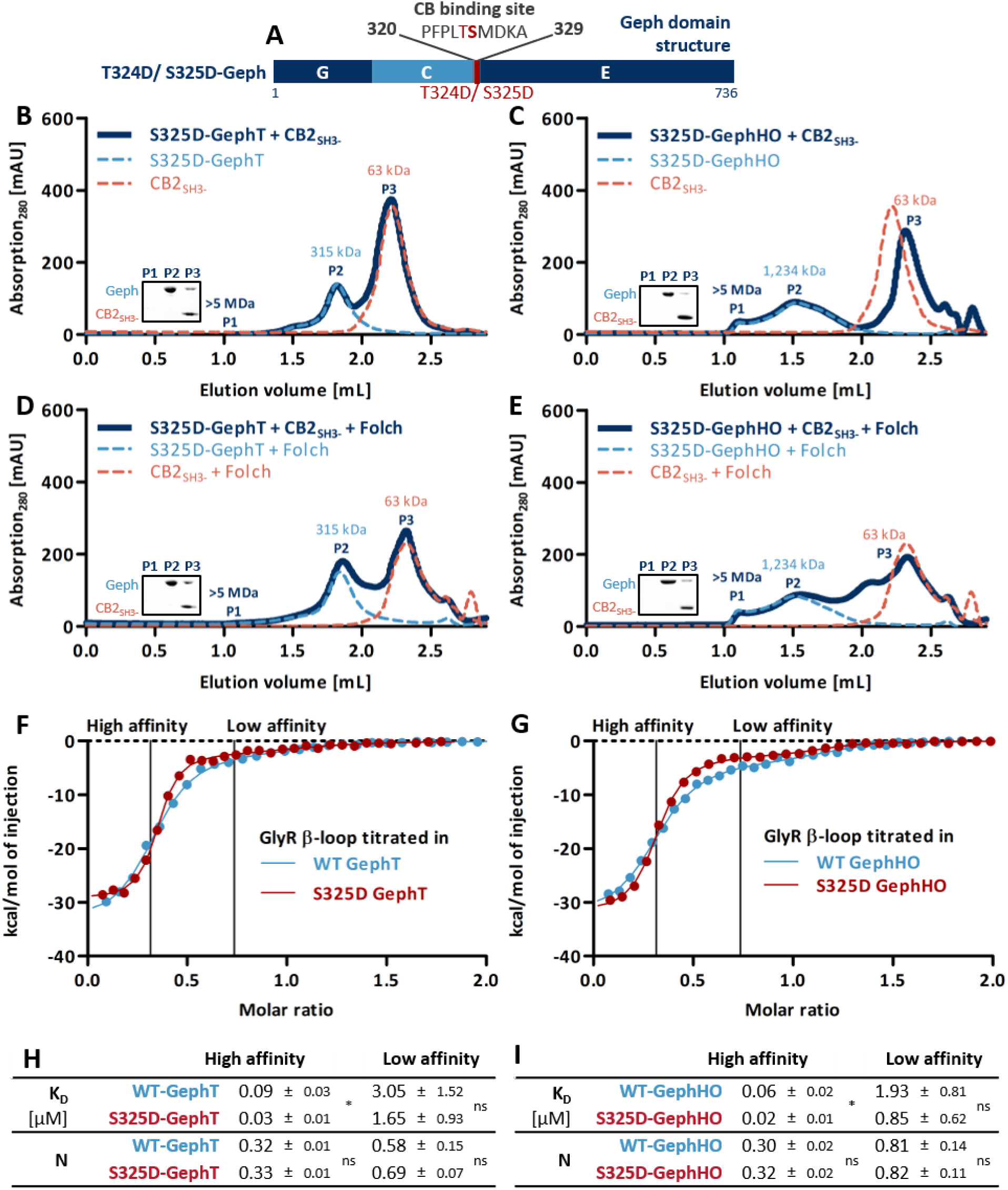
S325D-Geph does not form a complex with CB2_SH3-_ while the receptor binding ability is improved. ***A*,** Domain structure of the phosphomimicking mutant T324D-Geph and S325D-Geph with the phosphorylation sites highlighted in red. ***B – E,*** Representative SEC elution profiles of the different oligomeric states of S325D-Geph mixed with CB2_SH3-_ at equimolar ratios (dark blue line), alone or in the presence of Folch. Single Geph (dashed line, light blue) and CB2_SH3-_ (dashed line, orange), with or without Folch, serve as a reference within each panel. The MWs of the single proteins, determined according to standard protein calibration curve, are indicated. Insets depicting SDS-PAGE analysis of peak 1 (P1), peak 2 (P2) and peak 3 (P3) of the Geph-CB2_SH3-_ interaction run. ***F*, *G,*** Representative ITC binding isotherms of the GlyR β-loop titrated into the respective oligomeric state of WT-Geph or S325D-Geph. The binding data points were fitted using a two-side model (line) with the high and low affinity binding sites highlighted. ***H, I,*** Binding parameters derived from the ITC experiments including K_D_ (μM) and N (molar ratio). Results are expressed as mean ± SD (n = 4 from three independently purified protein batches) and were analyzed using unpaired, two-tailed Student’s t-test. For the high affinity binding site the K_D_ value was significantly decreased for both oligomeric states of S325D-Geph compared to WT-Geph (GephT: *p*=0.0113 (*); GephHO: *p*=0.0110 (*)). All other parameters were not significantly altered (ns = *p*>0.05).

Both phosphomimicking variants showed WT-like oligomerization behavior following recombinant expression and purification, indicating that Geph oligomerization is not affected by the amino acid exchanges (Figure S5A). Therefore, GephT and GephHO of the phosphomimicking variants were isolated and used for SEC interaction studies with CB2_SH3-_. T324D did not alter the Geph-CB2_SH3-_ complex formation (Figures S5C – S5F) and was therefore not subjected to further experiments. In contrast, in case of S325D-Geph no complex formation with CB2_SH3-_ could be observed regardless of the oligomeric state used (Figures 3B and 3C). Also the addition of Folch lipids could not recover the complex formation (Figures 3D and 3E). In conclusion, the phosphomimicking variant S325D within the CB binding site of Geph abolished the complex formation with CB2_SH3-_ suggesting a negative impact of phosphorylation at Ser325 on CB binding.

### S325D-Geph does not form membrane-associated microclusters upon co-expression with CB2_SH3-_ in HEK GPHN^−/−^ cells

Previous studies have demonstrated that co-expression of Geph together with CB, in its open conformation, relocates Geph from large intracellular aggregates (“blobs”) into small microclusters (diameter, 0.2 – 0.5 μm) located at the plasma membrane in human embryonic kidney (HEK) 293 cells (Kins et al., 2000). Given that S325D-Geph was not able to form a complex with CB2_SH3-_ within our SEC studies (Figures 3B – 3E), we investigated microcluster formation of S325D-Geph following co-expression with CB2_SH3-_ in HEK cells (Figure 4). To rule out that endogenous Geph does influence the clustering behavior of S325D-Geph, the interaction studies were performed in Geph knock-out HEK cells (HEK GPHN^−/−^, Figure S5G). Cells were transfected with mScarlet-IRES-mEGFP-Geph constructs, containing WT-Geph or S325D-Geph, either alone or together with moxBFP tagged CB2_SH3-_ (Figure 4A). Thereby, cells expressing mEGFP Geph were filled with mScarlet. As expected, WT-Geph as well as S325D-Geph alone did form large intracellular aggregates with no significant difference in cluster number and size between both Geph variants (Figures 4B and 4C). Upon co-expression with CB2_SH3-_ neither the number (Figure 4B) nor the size (Figure 4C) of the large intracellular S325D-Geph aggregates was altered, while in case of WT-Geph the expected formation of submembranous clusters could be observed (Figure 4A). For WT-Geph the cluster number was significantly increased (*p* < 0.0001, Figure 4B) while the cluster size was significantly decreased (*p* < 0.0001, Figure 4C) upon co-expression with CB2_SH3-_. Furthermore, the large intracellular S325D-Geph aggregates did not co-localize with CB2_SH3-_, which was diffusely distributed (Figure 4A). Thus, our results show that the negatively charged aspartate at position 325 of Geph abolishes the complex formation with CB2_SH3-_ and thereby prevents the membrane targeting effect of CB2_SH3-_ within mammalian cells.

**Figure 4.**
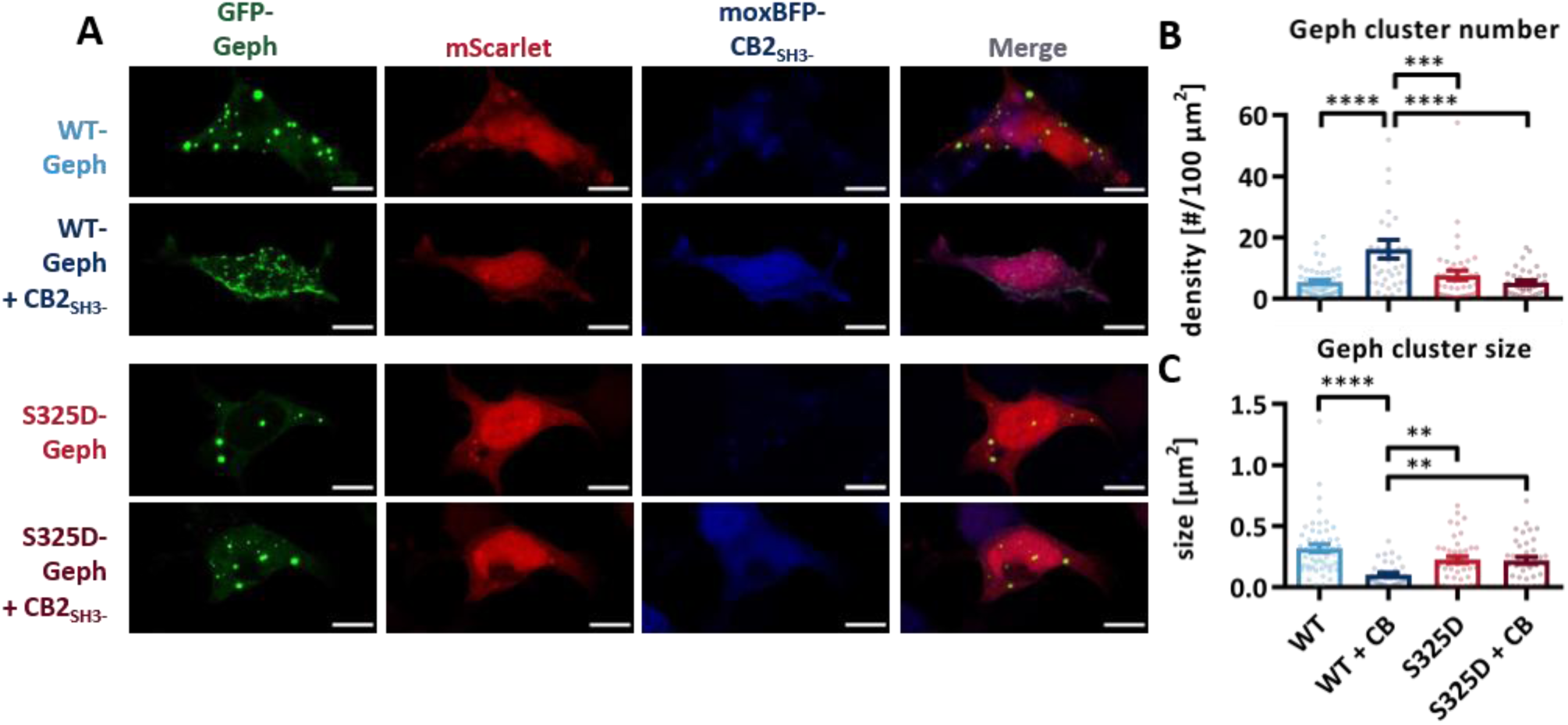
S325D-Geph does not form submembranous microclusters upon CB2_SH3-_ co-expression in HEK GPHN^−/−^ cells. ***A***, Representative confocal images of HEK GPHN^−/−^ cells co-expressing GFP tagged WT-Geph or S325D-Geph together with moxBFP-tagged CB2_SH3-_ and mScarlet as a cell filler (scale bars = 10 µm). S325D-Geph as well as WT-Geph form large cytoplasmic aggregates (‘blobs’) while CB2_SH3-_ induces the formation of WT-Geph microclusters at the plasma membrane. This microcluster formation is absent in case of S325D-Geph. ***B, C,*** Quantitative analysis of GFP-Geph clusters with individual data points and mean ± SEM displayed in the figures (n = 38 – 51 cells from three independent transfections). ***B,*** Quantification of the GFP-tagged Geph cluster number per 100 µm^2^ analyzed with Wilcoxon rank test (WT vs. WT+CB: *p*<0.0001 (****); WT+CB vs. S325D: *p*=0.00057 (***); WT+CB vs. S325D+CB: *p*<0.0001 (****); WT vs. S325D: *p*=0.646 (ns); WT vs. S325D+CB: *p*=0.994 (ns); S325D vs. S325D+CB: *p*=0.519 (ns)). ***C,*** Quantification of the mean GFP-Geph cluster size per cell [µm2] analyzed with Wilcoxon rank test (WT vs. WT+CB: *p*<0.0001 (****); WT+CB vs. S325D: *p*=0.002 (**); WT+CB vs. S325D+CB: *p*=0.005 (**); WT vs. S325D: *p*=0.034 (ns); WT vs. S325D+CB: *p*=0.032 (ns); S325D vs. S325D+CB: *p*=0.989 (ns)).

### S325D-Geph shows enzymatic activity and increased GlyR β-loop binding

To investigate whether the phosphomimicking variant S325D affects not only the complex formation with CB2_SH3-_ but also other functions of Geph, we investigated Moco synthesis and receptor peptide binding. Performing the previously described *in vitro* Moco assay (Belaidi & Schwarz, 2013), confirmed that the enzymatic activity of S325D-Geph was not altered (Figure S5H).

Receptor binding ability of S325D-Geph was investigated by isothermal titration calorimetry 378 – 426, GlyR β-loop), was titrated into the trimeric and high oligomeric state of S325D-Geph and WT-Geph (Figures 3F – 3I and Figure S6). Two binding sites were described for the interaction between Geph and the GlyR β-loop, displaying a high-affinity binding site in the sub-micromolar range and a low-affinity binding site with micromolar affinity (Grünewald et al., 2018; Herweg & Schwarz, 2012). Our ITC measurements revealed, that both oligomeric states of S325D-Geph were able to interact with the GlyR β-loop resulting in exothermic binding events revealing the low- and high-affinity binding site (Figures 3F and 3G). The stoichiometry of the formed complexes was comparable to WT-Geph in case of both binding sites (Figures 3H and 3I). Also the thermodynamic parameters of the interaction (binding enthalpy, entropy and free Gibbs energy) were not altered between S325D-Geph and WT-Geph (Figures S6E and S6F). However, in case of the high-affinity binding site, we observed a significantly increased, approximately three-fold higher, affinity of S325D-Geph compared to WT-Geph (Figures 3H and 3I, GephT: *p* = 0.0113, GephHO *p* = 0.0110). Thus, the ITC data shows that the phosphomimicking variant S325D exhibits an improved receptor binding.

### Impaired S325D-Geph clustering at GABAergic synapses in hippocampal neurons

Numerous studies have demonstrated the importance of CB in the formation and maintenance of Geph clusters at GABAergic synapses in selected regions of the mammalian forebrain, including the hippocampus (Papadopoulos et al., 2007, 2008; Tyagarajan, Ghosh, Harvey, et al., 2011). The phosphomimicking mutation S325D in Geph abolished complex formation with CB2_SH3-_ both *in vitro* (Figures 3B – 3E) and in HEK GPHN^−/−^ cells (Figure 4). Therefore, clustering of S325D-Geph was studied in murine hippocampal neurons using recombinant adeno-associated virus (rAAVs) based expression (Figure 5) (Liebsch et al., 2023). CamKII-positive glutamatergic cells were transfected with the recombinase Flpo and moxBFP as a cell filler. Flp-dependent mScarlet-WT-Geph and mScarlet-S325D-Geph constructs were expressed using rAAVs. The synaptic localization of Geph clusters was quantified using immunocytochemistry of the GABA_A_R ɣ2-subunit, as a marker of postsynaptic GABA_A_Rs, and the vesicular GABA transporter (vGAT), as a presynaptic marker (Figures 5C – 5H).

**Figure 5.**
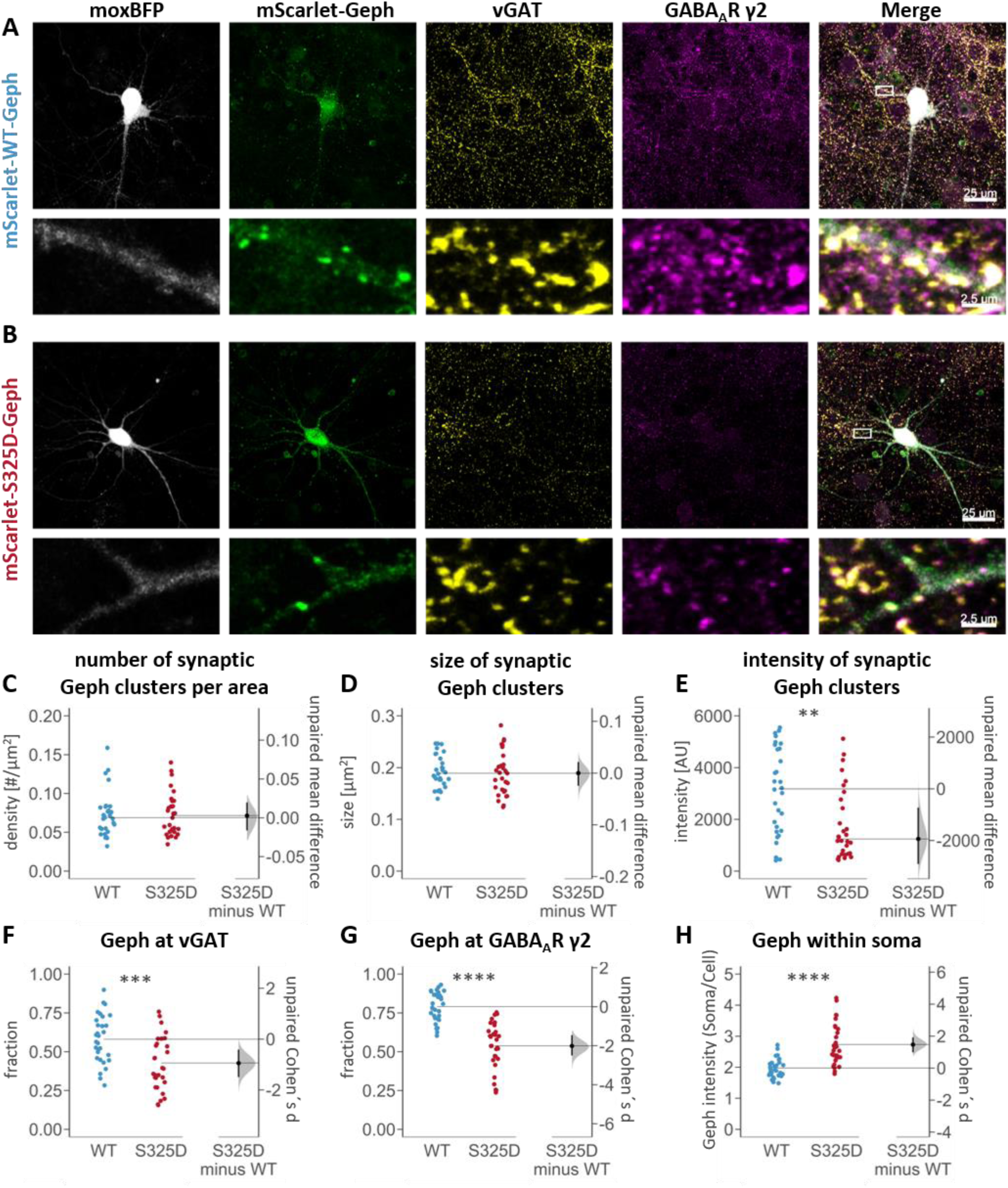
Impaired S325D-Geph clustering at GABAergic synapses in murine hippocampal neurons. Representative confocal images of hippocampal neurons expressing moxBFP-IRES-Flpo and mScarlet-tagged ***A,*** WT-Geph or ***B,*** S325D-Geph. ***C – H*,** Quantitative analysis of mScarlet-tagged Geph clusters was performed using an automated image analysis with the individual data points, mean, confidence intervals, and standard deviation displayed in the figures (n = 30 cells per condition from four independent cultures). ***C, D,*** Average number (per µm^2^) as well as average size (per cell) of synaptic clusters (vGAT-positive and GABA_A_Rɣ2-positive) is not different between both variants; Wilcoxon rank test *p* = 0.786 (ns) and *p* = 0.542 (ns), respectively. ***E,*** The average fluorescence intensity of synaptic S325D-Geph clusters (per cell) is significantly reduced compared to WT-Geph; Wilcoxon rank test *p* = 0.002 (**). ***F, G,*** The fractions of vGAT- and GABA_A_Rɣ2-positive clusters are significantly reduced in case of S325D-Geph; Student’s *t* test *p* = 0.0005 (***) and *p* = 4.1*10^-10^ (****), respectively. ***H,*** Ratio of somatic/whole cell mean Geph fluorescence intensity is significantly increased for S325D-Geph compared to WT-Geph; Student’s *t* test *p* = 1.2*10^-6^ (****).

WT-Geph and S325D-Geph formed clusters throughout dendrites and regularly co-localized with vGAT as well as the GABA_A_R ɣ2-subunit (Figures 5A and 5B). Density (Figure 5C) and size (Figure 5D) of synaptically localized S325D-Geph clusters were not altered. However, the intensity of synaptic S325D-Geph clusters was significantly reduced compared to WT-Geph (*p* = 0.002, Figure 5E). Furthermore, S325D-Geph displayed a significantly reduced proportion of synaptically localized clusters, with only 42.63 ± 17.22 % co-localized with vGAT compared to 58.12 ± 15.47 % for WT-Geph (*p* = 0.0005, Cohen’s d effect size d = −0.946, Figure 5F). The most drastic change, with the largest effect size, was observed in the proportion of Geph clusters co-localized with the GABA_A_R γ2 subunit, which was significantly reduced to 53.70 ± 15.06 % compared to 78.99 ± 9.43 % for WT-Geph (*p* ≤ 0.0001, Cohen’s d effect size d = −2.012, Figure 5G). These results demonstrate that clustering of S325D-Geph specifically at GABAergic synapses is impaired. Notably, clustering of the recombinantly expressed Geph variants was studied in the presence of endogenous Geph expressed in the neuronal cultures, suggesting that synaptic S325D-Geph clusters potentially arise due to oligomerization with endogenous Geph. Additionally, the somatic localization of S325D-Geph was significantly increased as the ratio of Geph intensity within the soma and the whole cell was increased to 2.73 ± 0.66 compared to 1.97 ± 0.30 for WT-Geph (*p* ≤ 0.0001, Cohen’s d effect size d = 1.470, Figure 5H).

Collectively, our cellular findings revealed a reduced association of S325D-Geph with GABAergic synapses alongside with an increased somatic localization within hippocampal neurons. This indicates that the abolished complex formation of S325D-Geph with CB, shown by SEC (Figures 3B – 3E) and HEK-293 cell studies (Figure 4), inhibits the recruitment and clustering of S325D-Geph at GABAergic synapses (Figure 5). These results highlight the importance of the high-molecular weight Geph-CB complex formation for proper GABAergic clustering and suggest a possible regulatory mechanism of Geph clustering at GABAergic synapses via phosphorylation at Geph Ser325.

## DISCUSSION

CB-induced formation of Geph clusters at GABAergic synapses is of fundamental importance, however, the molecular mechanisms underlying the Geph-CB complex formation remained poorly understood. Based on *in vitro* interaction and cellular studies using hippocampal neurons, we propose a model for the CB-dependent Geph clustering based on Geph oligomerization induced by CB binding. This process is facilitated by the interaction of CB with plasma membrane-resident PIPs and impaired by Geph Ser325 phosphorylation.

We discovered the formation of a high-molecular weight Geph-CB2_SH3-_ complex, implying that the complex formation is based on Geph self-oligomerization. Furthermore, variants of Geph, unable to form E-domain dimers (ddGeph), or lacking the trimerizing G-domain of Geph (GephE), were not able to form a high-molecular weight complex together with CB2_SH3-_. Those results led us conclude, that CB2_SH3-_ binding stabilizes the dimerized Geph E-domain and thereby together with G-domain trimers induces the formation of a Geph scaffold resulting in the high-molecular weight Geph-CB2_SH3-_ complex (Figures 1J and 1K).

A recent study, determining the affinity of the Geph-CB interaction using FRET measurements, revealed that a monomeric (dimerization-deficient) GephE variant displayed a much lower affinity (K_D_ = 44.1 µM) towards CB2_SH3+_ compared to dimerized GephE (K_D_ = 6.3 µM) (Imam et al., 2022). Their finding that Geph E-domain dimerization is not required for the initial binding event with CB but does enhance the affinity is in line with our conclusion, that CB binding induces the dimerization of the E-domain within full-length Geph. Potentially CB initially binds to the domain boundary of the C-domain and the monomerized E-domain, relieving the auto-inhibitory effect and thereby inducing E-domain dimerization, which in turn increases the affinity of the CB-Geph interaction.

An earlier study revealed that co-expression of Geph and CB2_SH3-_ in hippocampal neurons results in the formation of enlarged postsynaptic Geph clusters (0.81 ± 0.03 µm^2^), so-called “superclusters”, compared to Geph clusters formed without CB co-expression (0.24 ± 0.02 µm^2^) (Chiou et al., 2011). We propose that these “superclusters” are formed due to Geph network formation induced by CB2_SH3-_ and stabilized by PIPs at the postsynaptic membrane. Geph network formation based on domain self-oligomerization has been proposed for decades and was shown to be essential for Geph clustering in neuronal cells (Saiyed et al., 2007). Here we provide direct *in vitro* evidence for a CB- and PIP-dependent formation of such a Geph network.

Recent studies revealed that Geph network formation drives the formation of phase separated condensates, which is thought to be essential for compartmentalization of the inhibitory postsynaptic protein machinery into postsynaptic densities (Bai et al., 2021; Lee et al., 2024; Zhu et al., 2024). Thus, our finding, that CB induces Geph network formation stabilized by PIPs at the plasma-membrane potentially represents a molecular mechanism driving Geph condensate formation specifically at the postsynaptic membrane.

Recombinant Geph forms trimers (GephT) as well as distinct higher oligomeric states (GephHO). Our interaction studies revealed that the Geph-CB complex formation is dependent on the oligomeric state of Geph. While GephHO formed a high-molecular weight complex with CB2_SH3-_ alone, GephT only showed complex formation with CB2_SH3-_ in the presence of lipids. The mechanism and structural basis for GephHO formation is still unclear. However, we showed that E-domain dimerization is essential for the formation of GephHO as ddGeph, unable to form E-domain dimers, did not form any higher oligomers. Studies showing that the presence of the C-domain inhibits E-domain dimerization were solely performed with isolated domains or full-length GephT but not with GephHO (Bedet et al., 2006; Sander et al., 2013). Thus, it is possible that E-domain dimerization within GephHO is favored and therefore the affinity for CB is increased compared to GephT with monomerized E-domains.

An impaired formation of higher oligomers, as seen in the pathogenic G375D-Geph variant, impairs clustering at GABAergic synapses and led to epileptic encephalopathy, highlighting the importance of the formation of higher oligomers for synaptic Geph clustering (S. Kim et al., 2021). Indeed, for G375D-Geph, CB-induced submembranous microcluster formation in non-neuronal cells was reduced (Dejanovic et al., 2015), which further supports our finding, that the formation of higher Geph oligomers promotes the interaction with CB.

Our studies investigating the oligomeric state of native Geph from pig brain tissues, indicate that cytosolic Geph is mainly trimeric while membrane associated Geph is rather in an high-oligomeric state. Therefore, we conclude, that high-molecular weight complex formation with CB is preferentially formed by higher oligomeric Geph at submembranous sites. On the other hand, intracellularly Geph trimers with monomerized E-domains bind CB with low affinity and require additional plasma membrane-associated PIP interactions to induce Geph network formation (Figures 1J and 1K).

The ability of CB to bind PIPs is crucial for Geph targeting and clustering at CB-dependent GABAergic synapses (Harvey et al., 2004; Papadopoulos et al., 2017). Here, we show that PIPs do not only act as a membrane tethering component but do also stabilize the Geph-CB complex. Our finding that PIPs present at the plasma membrane, namely PI(3,4)P_2_, PI(4,5)P_2_ and PI(3,4,5)P_3_ (Ueda, 2014), promote the Geph-CB complex formation is in line the fact that those lipids are involved in the formation and maintenance of synapses (Dickson, 2019; Ueda, 2014; Volpatti et al., 2019). Consistently, PI(4,5)P_2_ was found to be required for CB-dependent Geph microcluster formation within non-neuronal cells (Kilisch et al., 2020). In aggregate, our results demonstate that the Geph-CB complex is stabilized and maintained by interaction with plasma-membrane-resident PIPs at postsynaptic sites (Papadopoulos et al., 2017).

Within a cryo-EM structure of a heteropentameric GABA_A_R, PI(4,5)P_2_ was identified to be bound to the cytosolic part of the α-subunits (Laverty et al., 2018). PI(4,5)P_2_ binding did not alter the channel function and thus was thought to modulate its synaptic localization (Laverty et al., 2018; Mennerick et al., 2014). Interestingly, residues mediating PI(4,5)P_2_ binding are only conserved in synaptic α-subunits (α1–3 and α5) and not in extrasynaptic subunits (α4 and α6) (Kasaragod & Schindelin, 2019). Collectively, we conclude that GABA_A_R-bound PI(4,5)P_2_ may recruit and stabilize the Geph-CB complex at synaptic sites thus facilitating the maintenance of Geph-CB-GABA_A_R clusters.

Besides the stabilization of Geph-CB clusters at postsynaptic membranes via PIPs, we also identified a regulatory mechanism impairing CB-induced Geph clustering via phosphorylation of Geph Ser325 within the CB binding motif. The CaMKIIα dependent phosphorylation of Geph Ser325 was identified within zebrafish mauthner cells upon auditory stimuli giving rise to increased intracellular Ca^2+^ levels activating CaMKIIα, which in turn led to an enhanced GlyR clustering (Ogino et al., 2019). Glycinergic inhibition plays a key role in rapid escape responses of zebrafish and thus Ogino and colleagues focused on the effect of Geph Ser325 phosphorylation on GlyR clustering (Ogino et al., 2019). CB is only required for Geph clustering at specific GABAergic synapses but dispensable at glycinergic synapses (Papadopoulos et al., 2007). Thus, the effect of Ser325 phosphorylation on CB-dependent Geph clustering at GABAergic synapses was not studied so far.

We have discovered that the inability of the phosphomimicking variant S325D-Geph to form a complex with CB, impaired its clustering at GABAergic synapses in hippocampal neurons. The increased somatic localization of S325D-Geph is in line with a previous study showing that Geph variants lacking the CB-binding motif were located within somatic aggregates upon expression in cortical neurons (Harvey et al., 2004). The impaired complex formation between CB and S325D-Geph potentially hampers CB-dependent transport via PI(3)P-containing sorting endosomes towards GABAergic synaptic sites leading to an increased somatic localization (Papadopoulos et al., 2017). In summary, these results strengthen the importance of the high-molecular weight Geph-CB complex formation for Geph-dependent GABAergic synapses formation and suggest an underlying regulatory mechanism via phosphorylation at Geph S325.

Our *in vitro* interaction studies characterizing the Geph-CB complex together with our studies in hippocampal neurons, collectively propose a model for the CB- and PIP-dependent Geph clustering at inhibitory postsynaptic sites (Figure 6). We suggest a CB-induced Geph oligomerization via E-domain dimerization that is stabilized at PIP-containing postsynaptic plasma membranes of GABAergic synapses and downregulated via phosphorylation of Geph Ser325.

**Figure 6.**
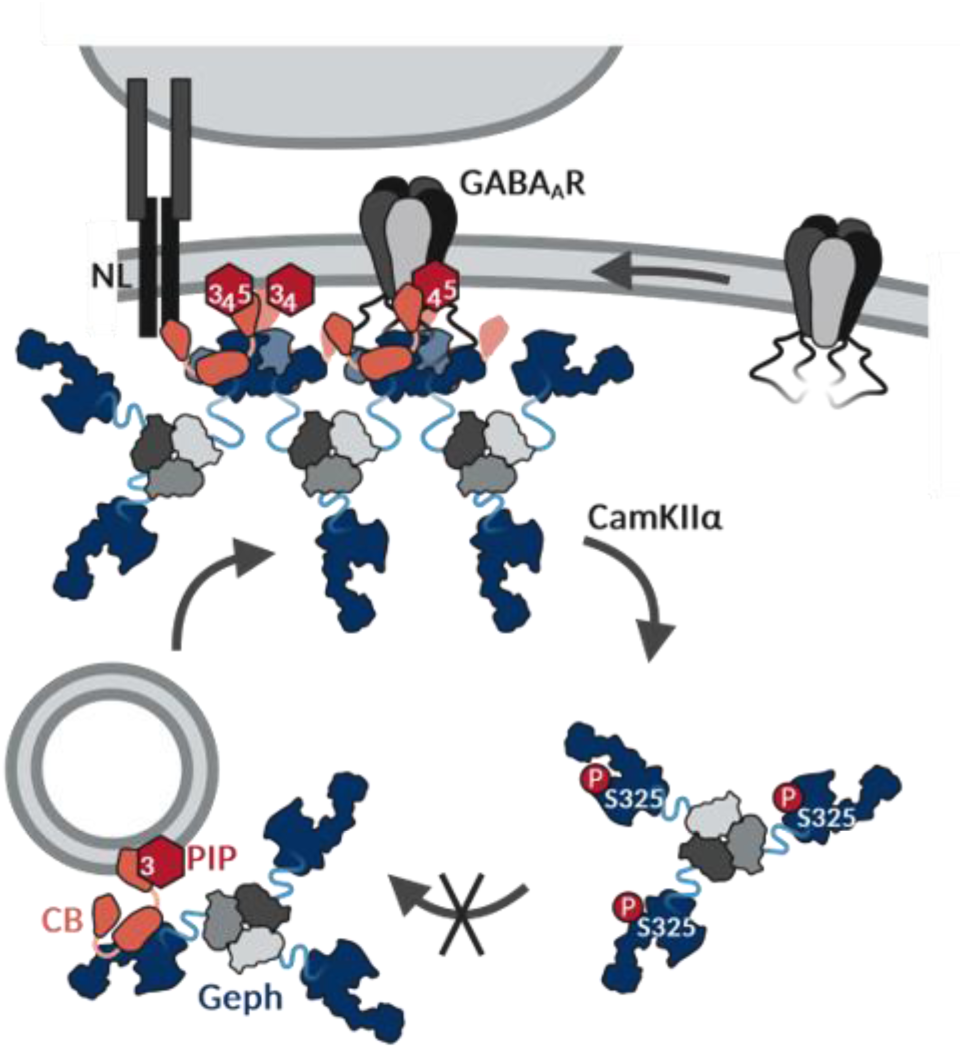
Proposed mechanism of Geph self-oligomerization induced by CB at PIP containing postsynaptic membranes regulated via Geph phosphorylation. ***A,*** Intracellularly Geph trimers with monomerized E-domains bind to CB with a low affinity. Either Geph itself (Imam et al., 2022) or other interactors such as TC10 (Kilisch et al., 2020) activate CB, so that it specifically binds PI(3)P at early/ sorting endosomes.(Papadopoulos et al., 2017) Prolonged interaction with TC10 induces a phospholipid affinity switch of CB towards plasma-membrane PIPs, resulting in the recruitment of the Geph-CB complex towards the plasma-membrane (Kilisch et al., 2020). ***B,*** Interaction of CB with plasma-membrane resident PIPs, PI(3,4,5)P_3_, PI(3,4)P_2_ and PI(4,5)P_2_, induces a conformational change within CB, that promotes the interaction with the dimerized Geph E-domain. Thereby, Geph E-domain dimerization is stabilized, which leads to the formation of a postsynaptic Geph network. Additionally, due to an unknown mechanism Geph associated with the plasma membrane adopts a higher oligomeric state (GephHO), which further promotes the complex formation with CB. PI(4,5)P_2_ bound to the cytosolic part of the GABA_A_R α-subunits can additionally stabilize this CB-Geph scaffold. ***C,*** The CB and PIP induced Geph network can be regulated via phosphorylation of Geph Ser325 via CaMKIIα activation upon increased neuronal activity (Ogino et al., 2019). This phosphorylation impairs the complex formation with CB, thereby leading to the removal of Geph from CB-Geph clusters at GABAergic synapses as well as an impaired transport via CB-PI(3)P containing endosomes. Figure was created with BioRender.com.

## ACKNOWLEDGEMENTS

We thank the Biocenter Imaging Facility and Dr. Matthias Gruhn for their support with confocal microscopy. We thank Prof. Dr. Andrea Musacchio and Dr. Raphael Gasper-Schönenbrücher for access to the mass photometer. Technical help from Monika Laurien and Simona Jansen is greatly appreciated. We thank Dr. Franziska Neuser and Julia Reich for their support with primary hippocampal neurons as well as Manuel Martinez-Osuna for his support with HEK GPHN^−/−^ cells. We thank Emanuel Bruckisch for his support in graphic design and illustration. Funding by the German Research Foundation is gratefully acknowledged by GS (RTG2550/1 project ID 411422114) and SP (SFB1430 - Project-ID 424228829). FB and SP were additionally funded by CMMC core funding (JRG XI). Support by CANTAR network funded by the Ministry of Culture and Science of the state of Northrine-Westphalia is acknowledged by SP.

## AUTHOR CONTRIBUTIONS

NB, FL, AM, JOG, PF, IR and FB performed the experiments; NB, SP and FL analyzed the data; NB, and GS wrote the manuscript; NB and GS designed the study.

## DECALARATION OF INTEREST

The authors declare no competing interests.

