## Supplementary Figures for "Phosphoinositide- and Collybistin-Dependent Synaptic Clustering of Gephyrin"

### SUPPORTING INFORMATION

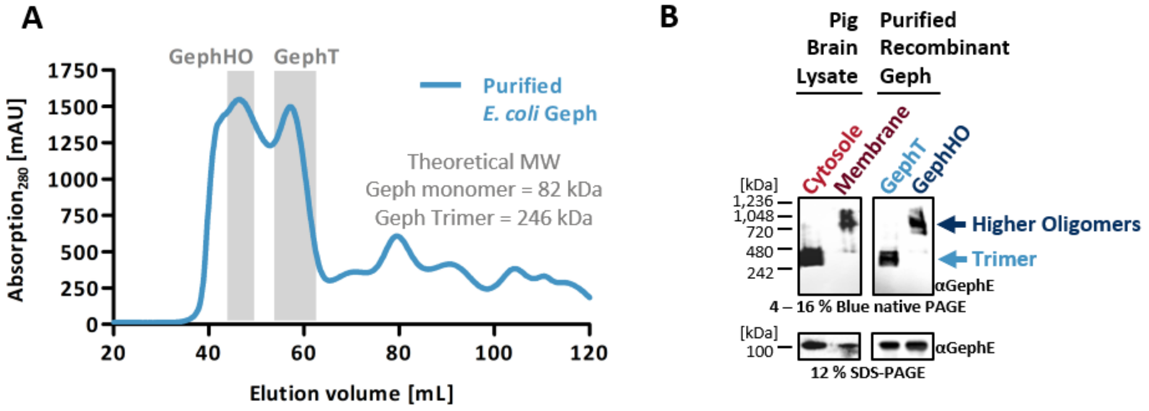


**Figure S1. Self-oligomerization of recombinant and native Geph into trimers and higher oligomers.**

***A,*** Preparative SEC elution profile of recombinant Geph expressed in *E. coli* after affinity purification with peaks corresponding to GephT and GephHO highlighted in grey. Theoretical MWs of monomeric and trimeric Geph are indicated. ***B,*** Blue native together with SDS-PAGE western blot analysis showing that native Geph from pig brain lysates forms trimers and higher oligomers comparable to recombinant Geph purified after expression in *E. coli*. Native brain lysates and recombinant Geph were loaded on the same gel, but recorded at different exposure times.

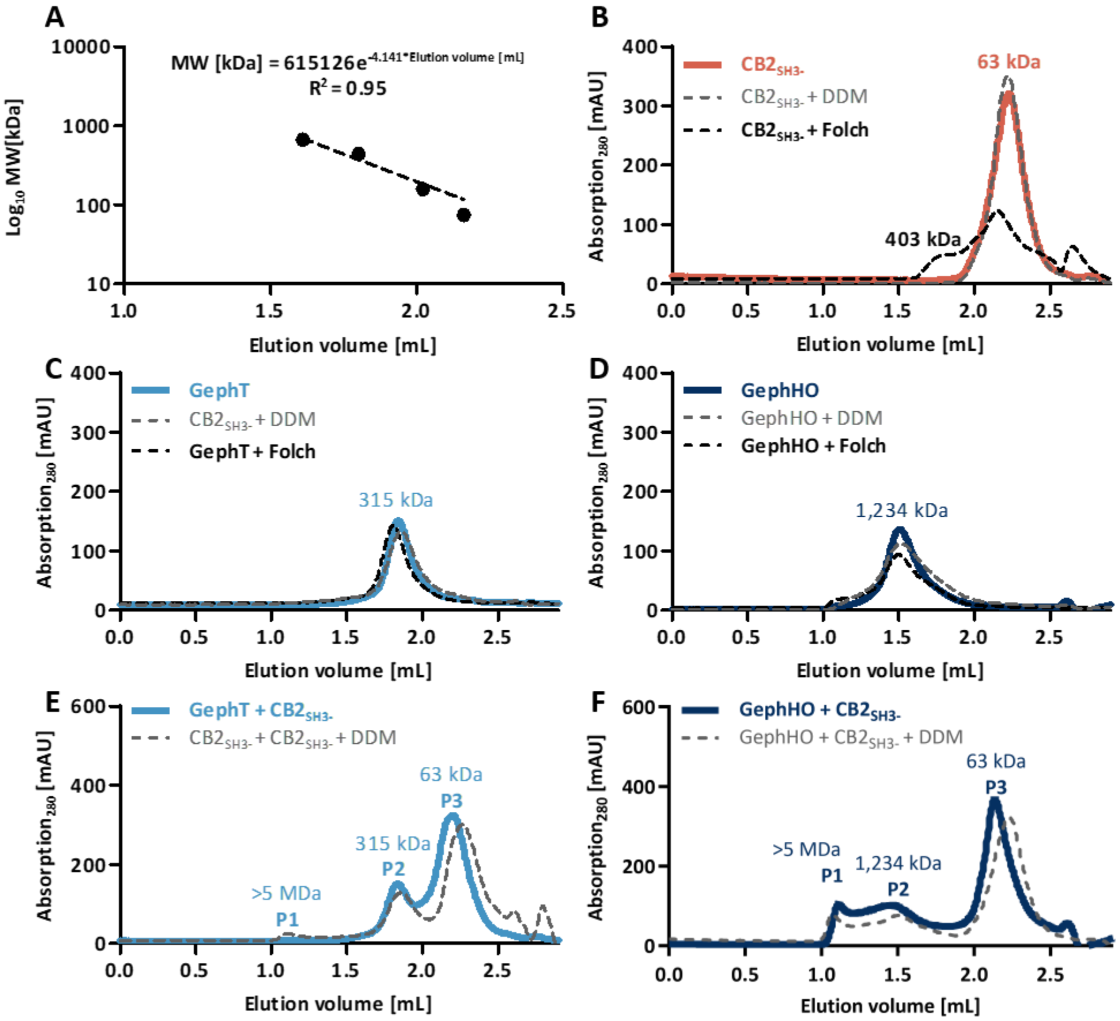


**Figure S2. SEC Interaction studies of Geph and CB2$_{SH3-}$ with Folch-DDM and DDM micelles.**

***A,*** SEC column (Superose 6 Increase 5/150 GL) calibration for the quantification of MWs using an exponential equation based on the elution volume of standard proteins (Conalbumin 75 kDa, Aldolase 158 kDa, Ferritin 440 kDa and Thyroglobulin 669 kDa). ***B,*** CB2$_{SH3-}$ interacts with Folch lipids within DDM micelles: Comparison of the SEC elution profiles of CB2$_{SH3-}$ in the presence or absence of Folch-DDM (shown in Figure 1), and in the presence of empty DDM micelles. ***C, D,*** Neither DDM nor Folch alter the self-oligomerization of Geph: Comparison of the SEC elution profiles of GephT or GephHO in the presence or absence of Folch-DDM (shown in Figure 1), and in the presence of empty DDM micelles. ***E, F,*** Addition of DDM does not affect the CB-Geph interaction: SEC elution profiles of the CB-Geph complex in the absence (shown in Figure 1) or presence or of empty DDM micelles.

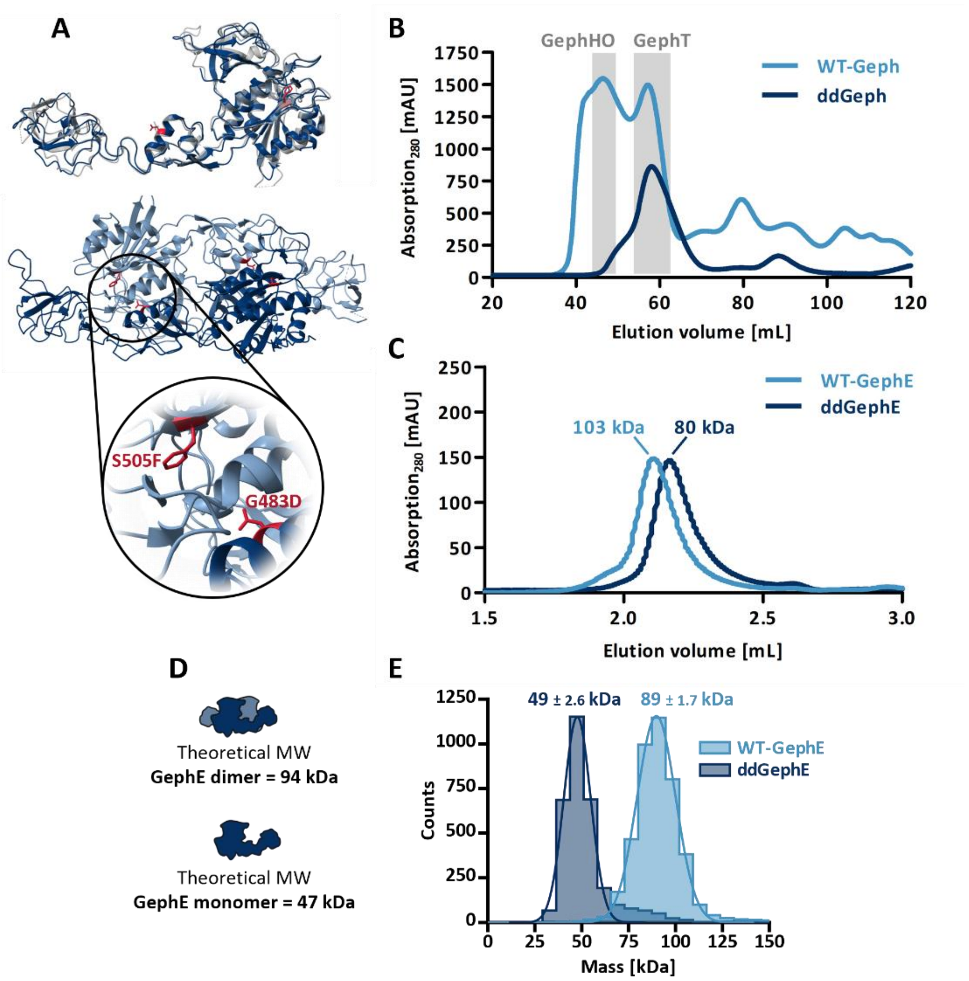


**Figure S3. The E-domain of ddGeph is not able to dimerize.**

***A,*** Top panel: Structural alignment of monomeric Cnx1 (grey, PDB: 5G2R) and the monomeric Geph E-domain (blue, PDB: 2FU3). The exchanged amino acids G483D and S505F (highlighted in red) are located in highly structurally conserved regions within both homologs. Bottom panel: Geph E-domain dimer (PDB: 2FU3) with the exchanged amino acids (highlighted in red) located at the dimerization interface (light blue vs. dark blue). The magnification depicts the close proximity of both amino acids to an α-helix, that is directly involved in the dimerization interface. The amino acid exchange from glycine to a larger, negatively charged aspartate in case of G483D possibly hampers the interaction with the adjacent α-helix of the second monomer. Due to the close proximity this α-helix is possibly also affected by the exchange from a serine to a larger, hydrophobic phenylalanine in case of S505F. ***B,*** Preparative SEC elution profile of full-length ddGeph directly after affinity purification, revealing that trimer formation is possible while the formation of higher oligomers is abolished. The elution profile of WT-Geph is shown as a reference. Peaks correlating to GephT and GephHO are highlighted in grey. ***C*,** SEC elution profile of ddGephE compared to WT-GephE, revealing that ddGephE elutes smaller than dimerized WT-GephE. MWs determined according to standard protein calibration curve are indicated. ***D*,** Theoretical MWs of the dimerized and monomeric Geph E-domain. Illustrations were created with BioRender.com. ***E*,** Representative mass distribution of ddGephE (dark blue) compared to WT-GephE (light blue) measured via mass photometry. The experimentally determined MWs are displayed in the figure as mean ± SD (n = 3 from three independent protein batches).

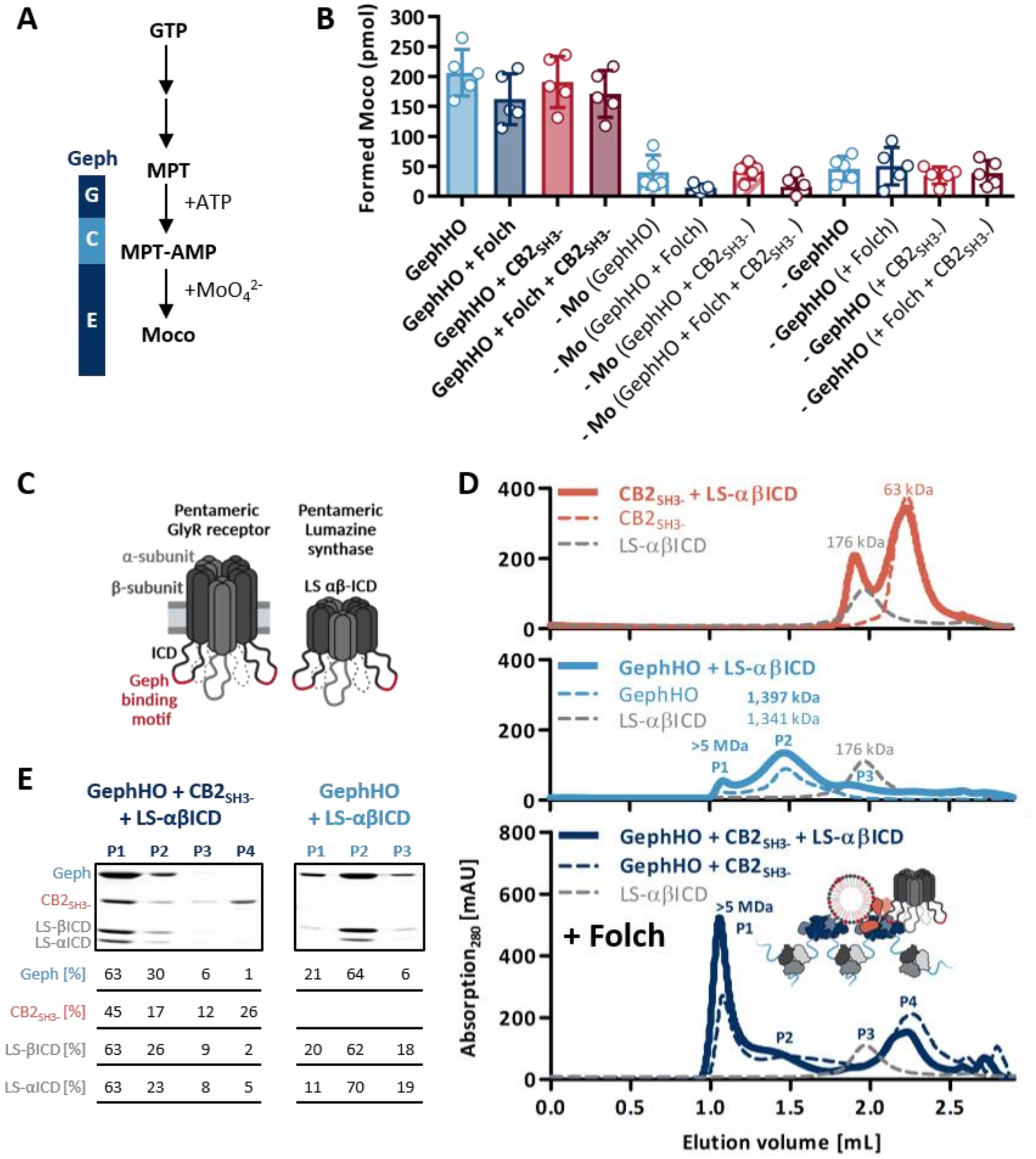


**Figure S4. Geph within the high-molecular weight Geph-CB2$_{SH3-}$ complex is active and can recruit the receptor model system LS-αβICD.**

***A,*** Moco biosynthesis pathway: Gephyrin G- and E- domain catalyze the last two steps from molybdopterin (MPT) to Moco. ***B,*** The Moco synthesis activity of Geph within the Geph-CB2$_{SH3-}$ complex is not impaired: *In vitro* Moco assay of the GephHO-CB2$_{SH3-}$ complex compared to GephHO alone in the presence or absence of Folch. Individual data points together with mean ± SD are displayed in the figure (n = 5 from two independently purified protein batches). Conditions containing Geph and Mo were analyzed using a 1way ANOVA, revealing no significant difference between GephHO alone or in complex with CB2$_{SH3-}$ in the presence or absence of Folch ($F(3, 16)=1.178$; $p=0.3493$; ns). Conditions without molybdenum (-Mo) or without Geph (-Geph) served as a negative control. ***C,*** Depiction of the lumazine synthase receptor model system with the incorporated ICDs of the GlyR α- and β-subunit (Macha et al., 2022). Figure was created with BioRender.com ***D,*** The >5MDa Geph-CB2$_{SH3-}$ complex is able to interact with the receptor model system LS-αβICD: SEC elution profiles of different combinations of GephHO, CB2$_{SH3-}$ and LS-αβICD mixed at equimolar ratios together with Folch (bold line). SEC elution profiles of the single proteins together with Folch (dashed lines) serve as a reference. The determined MWs, according to a standard protein calibration curve, of the single proteins as well as the protein complex are indicated. A scheme of the formed LS-αβICD-Geph-CB2$_{SH3-}$ complex together with Folch lipids is depicted in the bottom panel (created with BioRender.com). ***E,*** SDS-PAGE analysis of peak 1 (P1), peak 2 (P2), peak 3 (P3) and peak 4 (P4) of the respective SEC runs. Numbers within the table represent the relative band intensity [%] of Geph, CB2$_{SH3-}$ and both LS-αβICD subunits between P1, P2, P3 and P4.

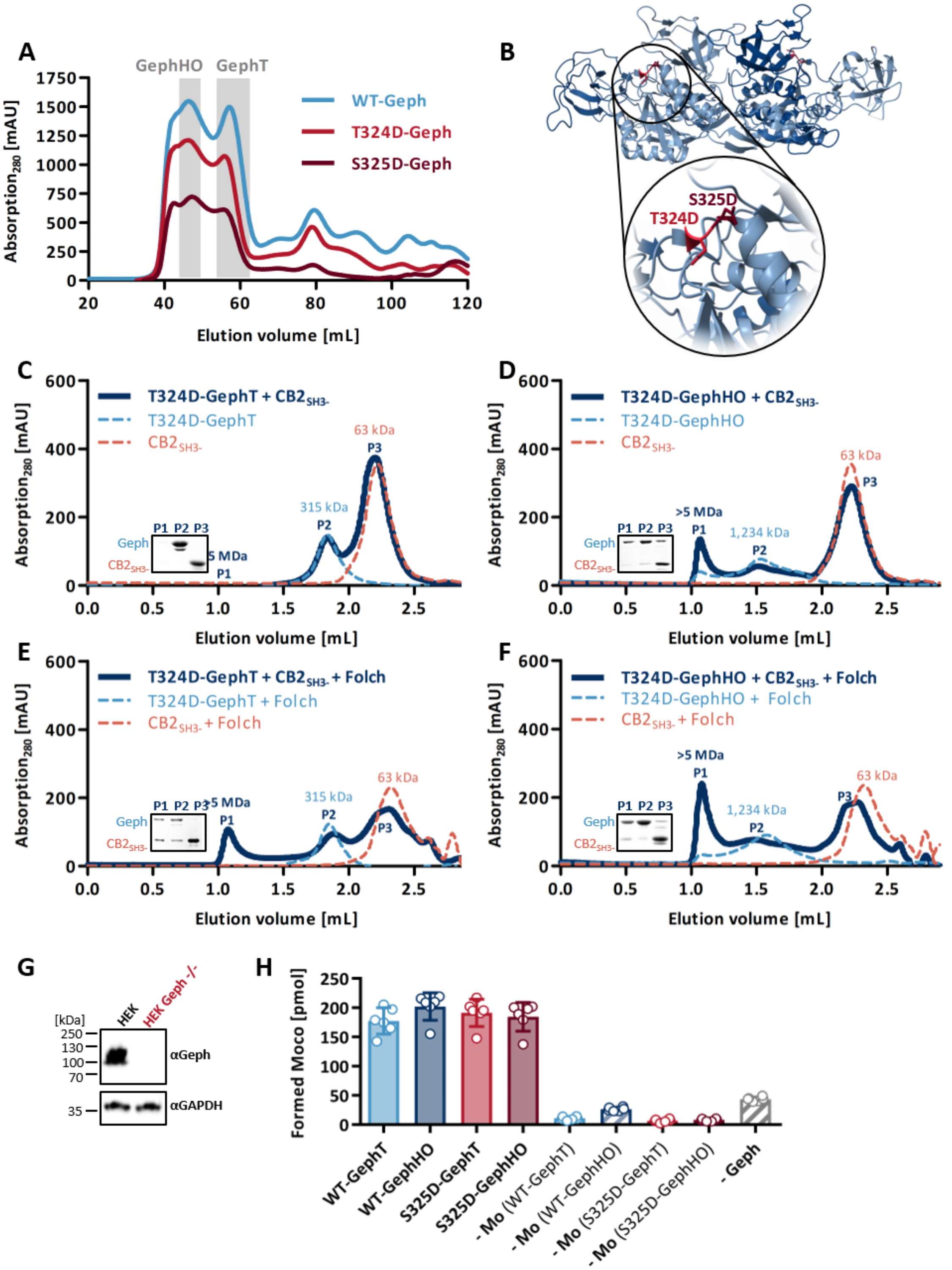


90

91 **Figure S5. Biochemical characterization of the phosphpomimicking Geph mutants,**

92 **T324D-Geph and S325D-Geph.**

***A,*** The phosphomimicking mutations do not impair the self-oligomerization into GephT and GephHO: Preparative SEC elution profile of T324D-Geph and S325D-Geph directly after affinity purification for separation of different oligomeric states. The elution profile of WT-Geph is shown as a reference. Peaks correlating to GephT and GephHO are highlighted in grey. ***B,*** Crystal structure of the Geph E-domain dimer (PDB: 2FU3) with the phosphomimicking mutations T324D and S325D highlighted in red. ***C – F,*** SEC elution profiles of T324D-Geph mixed with $CB2_{SH3-}$ at equimolar ratios (dark blue line), alone or in the presence of Folch. The determined MWs of the single proteins as well as the formed complex according to standard protein calibration curve are indicated. Single Geph (dashed line, light blue) and $CB2_{SH3-}$ (dashed line, orange), with or without Folch, serve as a reference within each graph. Insets depict SDS-PAGE analysis of peak 1 (P1), peak 2 (P2) and peak 3 (P3) of the Geph-$CB2_{SH3-}$ interaction run. ***G,*** Western blot analysis of HEK $GPHN^{-/-}$ cell lysates compared to HEK WT cell lysates, confirming that the Geph signal is absent for HEK $GPHN^{-/-}$ cells. ***H***, The enzymatic activity of purified S325D-Geph was measured using an *in vitro* Moco assay. The assay without Geph (−Geph) or molybdenum (−Mo) served as a negative control. Individual data points together with mean ± SD are displayed in the figure (n = 6 from two independently purified protein batches). No significant differences in Moco production between S325D-Geph and WT-Geph was observed, indicating that the enzymatic activity of S325D-Geph is not altered (1way ANOVA analysis: $F(3, 20)=1.19$; $p=0.3390$; ns).

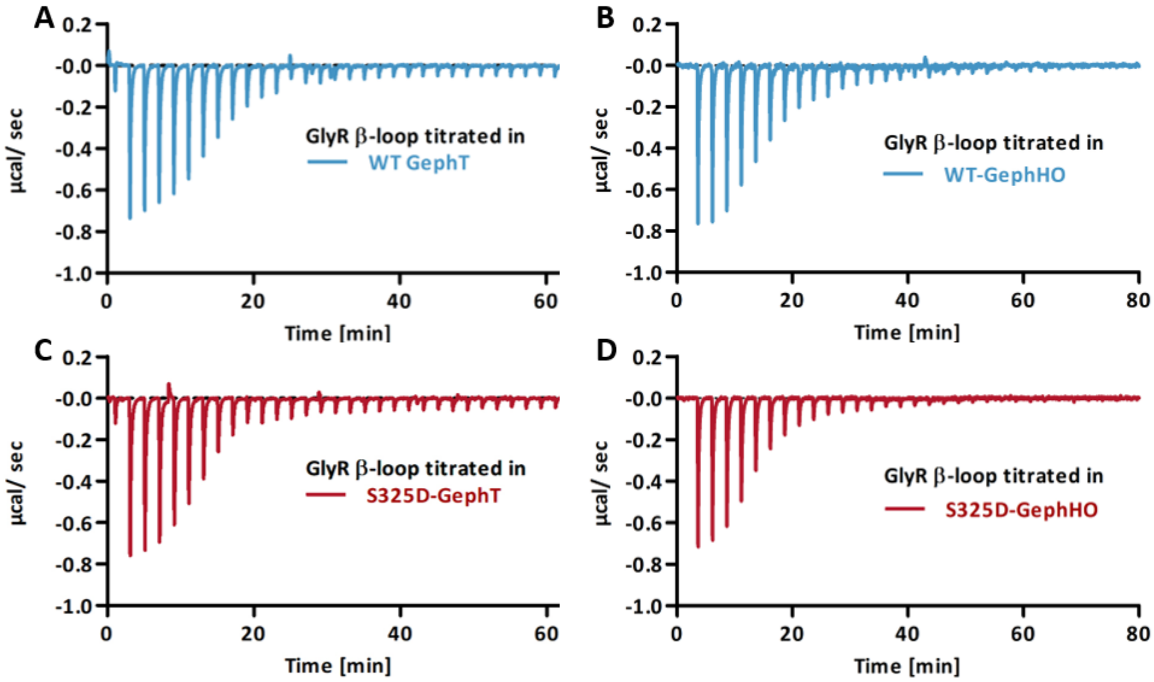


E

| | GephT | High affinity | | Low affinity | |
|---|---|---|---|---|---|
| $\Delta H$ | WT | -29.31 ± 4.31 | n.s. | -3.54 ± 2.24 | n.s. |
| [kcal/mol] | S325D | -32.16 ± 2.85 | | -4.25 ± 1.38 | |
| $-T\Delta S$ | WT | 19.29 ± 4.10 | n.s. | -4.37 ± 2.48 | n.s. |
| [kcal/mol] | S325D | 21.39 ± 2.97 | | -4.02 ± 1.36 | |
| $\Delta G$ | WT | -10.02 ± 0.27 | ** | -7.91 ± 0.40 | n.s. |
| [kcal/mol] | S325D | -10.77 ± 0.29 | | -8.27 ± 0.34 | |

F

| | GephHO | High affinity | | Low affinity | |
|---|---|---|---|---|---|
| $\Delta H$ | WT | -31.93 ± 1.55 | n.s. | -4.15 ± 0.72 | n.s. |
| [kcal/mol] | S325D | -33.51 ± 0.83 | | -3.49 ± 2.32 | |
| $-T\Delta S$ | WT | 21.66 ± 1.46 | n.s. | -4.02 ± 0.94 | n.s. |
| [kcal/mol] | S325D | 22.25 ± 1.21 | | -5.33 ± 2.79 | |
| $\Delta G$ | WT | -10.28 ± 0.22 | * | -8.16 ± 0.33 | n.s. |
| [kcal/mol] | S325D | -11.26 ± 0.60 | | -8.82 ± 0.68 | |

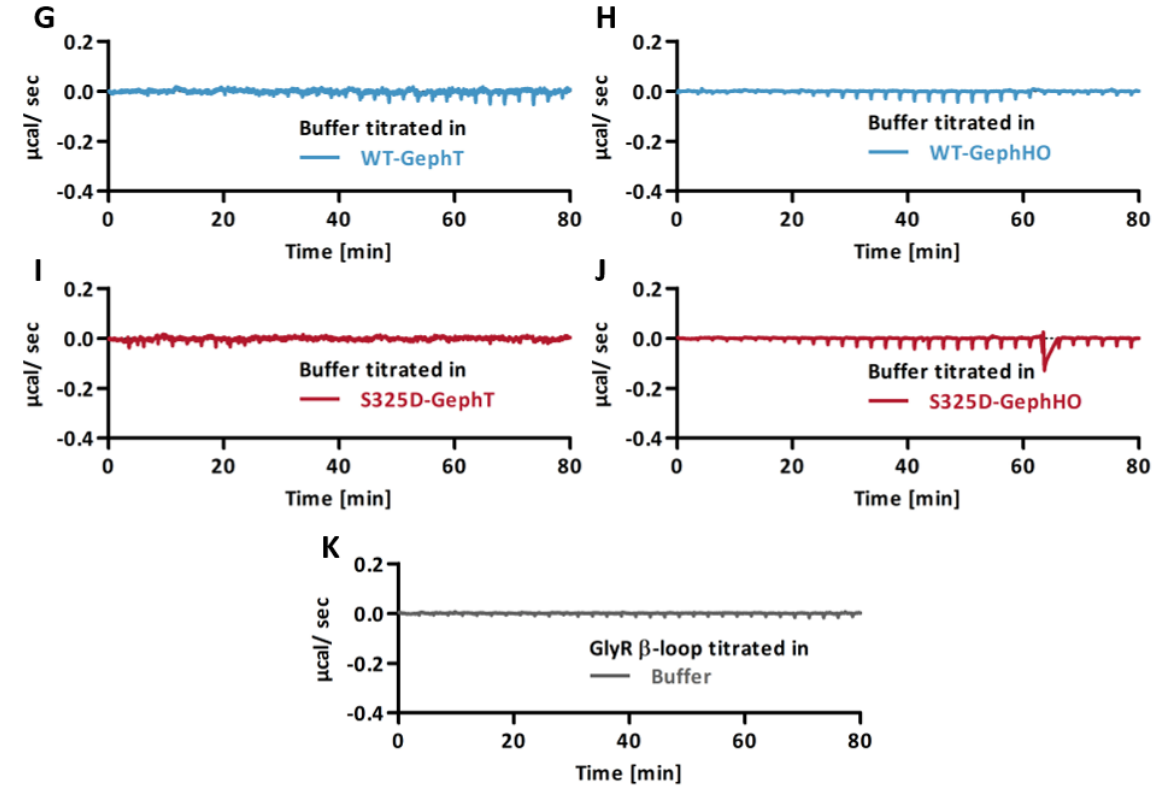


**Figure S6. S325D-Geph receptor binding ability measured via ITC.**

***A – D*,** Representative ITC thermograms, of the GlyR β-loop titrated into the respective oligomeric states of WT-Geph (blue) and S325D-Geph (red). ***E, F,*** Thermodynamic parameters derived from the fitted ITC experiments including binding enthalpy ΔH (kcal/mol), binding entropy −ΔTS (kcal/mol) and free Gibbs energy ΔG (kcal/mol). Results are expressed as mean ± SD (n = 4 from three independently purified protein batches) and were analyzed comparing WT-Geph and S325D-Geph of the respective oligomeric state using Student's t-test. In case of the high affinity binding site, binding enthalpy (H) and entropy (-TΔS) resulted in an energetically more favorable interaction with a significantly lower free Gibbs energy in case of S325D-Geph compared to WT-Geph (GephT: $p$ = 0.0091 (**); GephHO: $p$ = 0.0225 (*)). Between all other parameters no significant difference was observed (n.s. = $p$>0.05). ***G – K***, ITC thermograms of buffer titrated into the respective Geph variants or the GlyR β-loop titrated into buffer reveal that there are no unspecific binding events detected by the isolated proteins.

***References***

Macha, A., Grünewald, N., Havarushka, N., Burdina, N., Nagel-steger, L., Niefind, K., & Schwarz, G. (2022). Pentameric assembly of glycine receptor intracellular domains provides insights into gephyrin clustering. *BioRxiv*. https://doi.org/10.1101/2022.11.10.512828
